# A map of Non-translated RNA (nt-RNA) junctions in cancer genomes: a database resource of unproductive splicing

**DOI:** 10.1101/2025.06.15.659434

**Authors:** Dan Huang, Tsz-Ki Kwan, Suk-Ling Ma, Nelson LS Tang

## Abstract

**Background:** Non-translated transcripts (nt-RNAs) with frame-shifts or premature termination codons resulting from alternative splicing events (ASE), have been recently found at unexpectedly abundant in transcriptomes of cancer tissue. However, their full genomic spectrum has not yet been fully elucidated. This study comprehensively characterised the expression of signature junctions of these nt-RNA (termed “toxic junctions” here) of both known and novel nt-RNA across multiple cancer types and investigated their potential as biomarkers.

**Methods:** RNA-seq data of ∼6,000 samples, including the tumor and normal samples for 13 cancer types were retrieved from The Cancer Genome Atlas database (TCGA) together with data from Cancer Cell Line Encyclopedia (CCLE) project. Due to the difficulty in quantifying the entire transcript isoform of nt-RNA, we pioneered an algorithm to focus exclusively on the expression of junctional reads, which also circumvented the limitation of non-directional RNA-seq of TCGA data. We showed that the majority of nt-RNA is associated with at least one toxic junction. We built a comprehensive catalogue of known nt-RNA toxic junctions from genome databases. And novel toxic junctions were also identified by a new junction-focused algorithm from the higher quality discovery subsets of TCGA data. Splicing in Ratio (SiR) was used to quantify ASE leading to nt-RNA, enabling:

Differential expression analysis between cancer and normal tissue and across cancer types.

Identification of different profiles of nt-RNA abundance and various factor which may be the causes of differential nt-RNA abundance and SiR results

Identification of specific nt-RNA and toxic junctions that were expressed in various cancer (and/or normal tissue) types.

Assessment of nt-RNA and their toxic junction expression as biomarkers or prognosis indicators.

**Results:** We profiled the expressed known nt-RNA (toxic) junctions of known transcripts and discovered ∼22,000 novel toxic junctions out of ∼250,000 novel junctions found in the transcriptome data. The expression of nt-RNA was as high as 10% of all transcripts of the corresponding gene in cancer transcriptomes. Interestingly, some signature toxic junctions of nt-RNA are expressed in even higher quantities, e.g. up to 50% or more, which is reminiscent of a heterozygous mutation. We identified distinct patterns between cancer and normal samples, including example of nt-RNA expressing toxic junctions exclusively in normal or tumor samples. Clinically relevant examples included ANXA6 in breast cancer, where the nt-RNA isoform showed significantly higher expression in tumors (p=1.8e-15). In kidney renal clear cell carcinoma (KIRC), a significant isoform switch of ESYT2 based on the RNA-seq data was confirmed. The Kaplan-Meier survival curves showed that samples with the higher expression ratio of ESYT2-L are associated with better survival (p=2.0e-06). Unsupervised clustering showed that SiR results of 150 toxic signatures defined 4 subgroups of patients with different prognosis. Through principal component analysis (PCA), PC1 and PC2 can be used as an independent prognosis biomarkers. nt-RNA accounting for these PCs included splicing factors SRSF3 and CLK1, where CLK1 phosphorylates SRSF3 to promote exon 4 inclusion in both genes.

**Conclusions:** In summary, the expression profiles of all known and novel toxic junctions were explored using pan-cancer RNA-seq data. A dual 10% rule emerged from this study: ∼10% of novel junctions were toxic junctions associated with nt-RNA, and up to 10% of RNA transcripts inside a cell were also nt-RNA. The SiR metric enables accurate quantification of unproductive splicing and identification of cancer biomarkers. Our findings reveal that unproductive splicing represents functionally important post-transcriptional regulation in cancer. These expression profiles allow researchers to study the expression of nt-RNA signature junctions or novel signature junctions in or near the genes they are interested in, which could provide a new direction for their research. The SRSF3-CLK1 regulatory mechanism provides insights into splicing dysregulation. Our comprehensive toxic junction catalogue serves as a valuable resource, suggesting that targeting unproductive splicing pathways may offer novel therapeutic strategies for cancer treatment.

**Data availability:** The catalogue is available on GitHub and UCSC browser. https://github.com/danhuang0909/nt_database for GitHub overview https://genome.ucsc.edu/s/dandan_0909/hg38_all_new_nr for genome browsing of all novel (unannotated) toxic junctions https://genome.ucsc.edu/s/dandan_0909/hg38_5_26 for toxic junctions in known (annotated) nt-RNA.

## 1. Introduction

With the advancement of deep sequencing technologies, the complexity of the human transcriptome has been unveiled. The paradox between the small number (∼20,000) of protein-coding genes and the organismal complexity is intriguing. It indicates that many regulatory or complex mechanisms are involved in shaping the functional elements of human cells. One contributing mechanism is alternative splicing: a gene may produce multiple mRNAs transcript isoforms through various RNA processing mechanisms. Previous studies based on high-throughput sequencing show that more than 90% of genes undergo alternative splicing which largely increases the diversity of protein isoforms (Wang et al. 2008; Pan et al. 2008).

A commonly reported alternative splicing event (ASE) is exon skipping, in which a whole exon is either included or excluded in the mature RNA. In fact, alternative splicing can arise from at least four other basic mechanisms: alternative 5’ donor sites, alternative 3’ acceptor sites, mutually exclusive exons and intron retention (Black 2003; Pan et al. 2008). Alternative splicing may have very different effects on the structure of proteins, ranging from completely benign to completely defective (loss-of-function). For example, an inclusion or exclusion of functional motifs due to inclusion or loss of amino acids may result in changes in protein function. Also, the most severe functional alteration due to ASE may lead to production of frame-shifts transcripts which are then degraded by nonsense-mediated decay (NMD). therefore, the target protein is not produced at all. Such splicing errors can have profound consequences on protein function if tumor suppressor genes or oncogenes are involved. Such wasteful production of NMD targeted mRNA transcript was believed to be cleared up by rapid degradation. However, recent large-scale studies revealed that they were found in the steady-state mRNA transcript pool inside cancer cells and even in normal cells, though at a much lower quantities (Kahles et al. 2018).

The selection of the splicing sites was regulated by the trans-acting regulatory factors (various RNA binding proteins and splicing factors) that bind to the cis-acting RNA sequence elements (binding sites) of the pre-mRNA. The trans-acting regulatory factors include serine/arginine (SR)-rich proteins, heterogeneous ribonucleoproteins (hnRNPs) and some other unknown factors. The cis-regulatory sequence elements are short sequence motifs, usually consisting of 6-10 nucleotide bps. They could be divided into four classes, according to their position and functions: exon splicing enhancer (ESE), intron splicing enhancer (ISE), exon splicing silencer (ESS) and intron splicing silencer (ISS). SR proteins could recognize and bind to the splicing enhancers (ESE or ISE) and then enhance the recruitment of the spliceosome to enhance splicing. On the contrary, hnRNP proteins, which repress splicing by the assembly of the adjacent splice site and the spliceosome by binding to splicing silencers, bind to ESS or ISS (Smith and Valcárcel 2000; Wang and Burge 2008) Aberrant splicing, including unproductive splicing, in cancer is frequently driven by dysregulation of these splicing factors. Recurrent mutations in genes encoding core spliceosomal proteins, such as SF3B1, SRSF2, U2AF1, and ZRSR2, are commonly observed in various cancers, particularly in haematological malignancies. These mutations can alter the specificity of the spliceosome, leading to the selection of alternative splice sites and the production of unproductive transcripts. Furthermore, changes in the expression levels of splicing factors, either through upregulation or downregulation, can disrupt the normal equilibrium of splicing regulation, promoting unproductive splicing events like the inclusion or skipping of poison exons. The frequent occurrence of mutations and expression changes in splicing factors within cancer cells underscores their critical role in maintaining splicing fidelity and their significant contribution to the widespread splicing dysregulation observed in tumors.

Cryptic splice site activation is another mechanism contributing to unproductive splicing. Mutations can occur to activate splice sites within pre-mRNAs that are normally silent. This can result in the inclusion of intronic sequences or the skipping of exonic sequences, often leading to the generation of non-functional or truncated proteins. For instance, mutations in the SF3B1 gene can alter the recognition of the branch point sequence, leading to the use of cryptic 3’ splice sites. The activation of these normally ignored splice sites underscores the sensitivity of the splicing machinery to even subtle genetic changes, which can have far-reaching effects on the transcriptome.

In most biomarker or drug-discovery studies, gene expression profiles are used to study the mechanism of disease development. In the past, it was done typically using microarray to obtain a summary expression value for every gene. However, genes may have multiple isoforms which may have different functions. Genes may show differential expression level at isoform level but not gene level. With the development of sequencing technology, quantification of isoforms in different samples are possible. In recent years, the regulatory mechanism of genes in diseases at isoform-level attracts more and more interest. Zhang et al. demonstrated that isoform-level data have a better performance in distinguishing cancer samples from normal samples than gene-level data (Zhang et al. 2013). The process of splicing isoform switch is widely present in different cancers, indicating that alternative splicing plays a vital role in the development of disease (Zhao et al. 2016; Vitting-Seerup and Sandelin 2017).

Unproductive splicing is increasingly recognised as a significant contributor to the initiation and progression of various cancers. The ability of unproductive splicing to modulate gene expression and protein isoforms allows it to influence fundamental cellular processes that are often dysregulated in cancer. One key mechanism is the inactivation of tumor suppressor genes. By introducing premature termination codons into the transcripts of these genes and triggering their degradation via NMD, unproductive splicing can effectively lead to a loss of function, which is a hallmark of cancer initiation. Several examples illustrate this point, including the mis-splicing of crucial tumor suppressor genes like BRCA1 and PTEN in breast cancer, as well as KRAS in lung cancer, all of which have been reported to promote the onset of tumorigenesis.

RNA-seq is a technique for quantifying the transcriptome based on second-generation sequencing, and it can be used to identify gene sequences of previously unknown mRNA transcripts produced by ASE. RNA-seq could discover and quantify transcript at genome scale with features such as more accurate quantification, higher reproducibility, more extensive detection range, lower cost and more reliable analysis (Wang et al. 2009). It is a powerful tool for the in-depth study of the transcriptome.

The advent of large-scale genomic datasets, such as those generated by The Cancer Genome Atlas (TCGA) and the Genotype-Tissue Expression (GTEx) project, has provided unprecedented opportunities to investigate the landscape of splicing alterations in cancer, including unproductive splicing events. TCGA, a comprehensive initiative, has amassed genomic, transcriptomic, and proteomic data from a vast array of over 30 different cancer types, encompassing thousands of patients. Several studies have leveraged these rich datasets to specifically investigate unproductive splicing events across a multitude of cancer types. A significant pan-cancer analysis utilising TCGA data revealed a widespread increase in alternative splicing events in tumor samples when compared to their normal counterparts. This study demonstrated that, on average, tumors exhibit approximately 30% more alternative splicing events than normal tissues, with some cancer types showing even greater increases (Kahles et al. 2018).

However, transcriptome reconstruction is extraordinarily complex and challenging, particularly with short and non-stranded RNA sequencing which was used by TCGA. Firstly, the gene expression varies widely, some genes such as housekeeping genes may have thousands of supported reads, while others may only be supported by several reads. Also, a gene may express multiple transcripts, and the similarity between transcripts is high. Another problem is that the mature RNA, incompletely spliced precursor RNA, and the RNA being decayed exist at the same time in the cell, which leads to different reads coverage in different regions of a transcript. These limitations make it very difficult to determine a sequencing read to a particular mRNA transcripts isoform. Previous attempts to catalog ASE in cancer or normal cell genomes are based on mRNA transcripts. On the other hand, we took a junction-centric approach, we confined our analysis to only (exon) junctional reads. Junctions are the boundary of 2 adjacent exons which are joined together by splicing of the pre-mRNA. From there, we can identify toxic junctions which cause frame-shift of the protein coding sequence (CDS) with high degree of confidence. These junctions will result in unproductive transcripts (nt-RNA).

This report covers three aspects of study of nt-RNA.

1. A genome-wide catalog of nt-RNA junctions.
2. Regulation of expression of nt-RNAs and some examples in cancer genomes.
3. Genes in the splicing machinery with their nt-RNAs form a network in determining the prognosis of kidney cancer.

Although there are many catalogues of ASE which dated back to before the era of massive parallel sequencing, e.g. AceView (Thierry-Mieg and Thierry-Mieg 2006), these earlier dataset are limited in scope. More recently established databases tried to collect all ASE which are too plenty making functional annotation was not possible for most of the ASE (Kahles et al. 2018). Therefore, other researchers tried to develop another type of databases to understand the origin of ASE by identifying the causative DNA variants in cis (Cotto et al. 2023). Last year, Fair et al reported a database combining both causative DNA variant (as a QTL) and a key functional sequel of ASE that is unproductive splicing using nascent RNA-seq in lymphoblastoid cell lines of the HapMap project (Fair et al. 2024). Unproductive mRNA transcripts accounted for up to 15% of the steady-state mRNA pool in these cell lines. However, the genomes of these cell lines are considered to be germline, there was no large-scale survey of scale and property of ASE in cancer genomes focusing on unproductive splicing. In this paper, we focused on a specific signature, **toxic junctions,** to unambiguously identify unproductive transcripts. This is different from most other databases which were developed from exon-centric models. It is a first large-scale attempt to perform a junction-centric search for non-translated mRNA (nt- RNA) in the cancer genome. We focused on only this type of ASE because the loss-of-function sequel of nt-RNA can be sure. Please see Figure 1. Resources are also provided in Github and UCSC genome browser for validation and exploration by other researchers.

**Figure 1.**
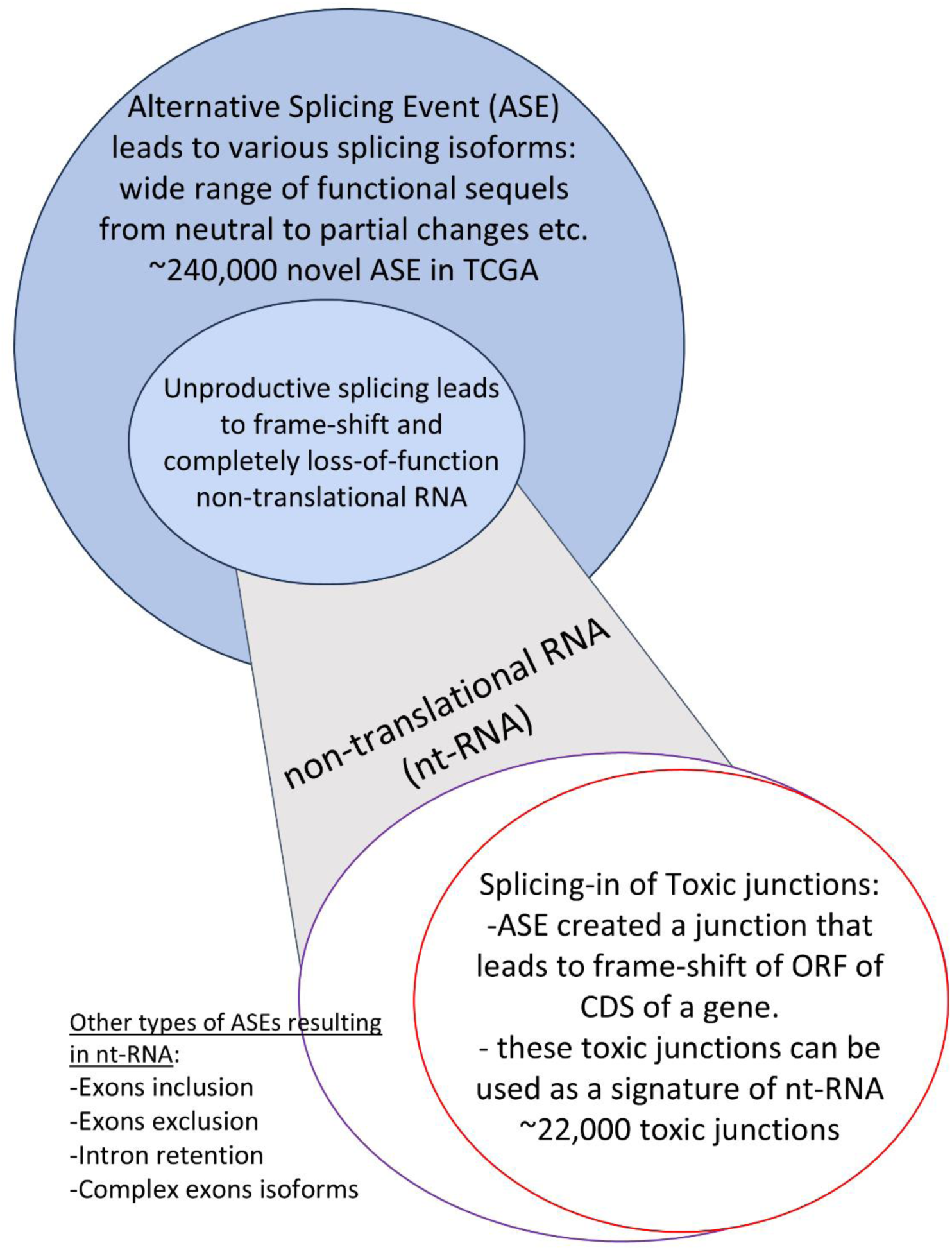
Identification of toxic junction. Previous works and databases focused on the exons as the functional unit of splicing (exon-centric). We carried out the first large-scale junction-centric search for non-translated mRNA in the cancer genome. We confined our analysis to only junctional reads of RNA-seq. Junctions are the boundaries of two adjacent exons that are joined together by the splicing of the pre-mRNA. From there, we can identify toxic junctions that cause a frame shift in the protein-coding sequence (CDS) with a high degree of confidence. These toxic junctions will result in non-translated transcripts (nt-RNA) and they can be used as splicing signatures for further analysis of cancer transcriptome.

## 2 Materials and Methods

### 2.1 Overview of analysis workflow to identify and evaluate toxic junctions of nt-RNA

The focus of this study is on a class of transcript isoforms that are defective in terms of protein translation. We called them non-translated transcripts (nt-RNA). They are isoform transcripts produced from the protein-coding gene but carrying a frame-shifted CDS or a premature stop codon, usually due to alternative splicing. Most nt-RNA have unique exon-exon splicing junctions within that nt-RNA. Therefore, these toxic splicing junctions can be used as signatures to identify unambiguously nt-RNA transcripts in RNA-seq reads.

Contrary to previous understanding, ASE are abundant in cancer transcriptomes and even in normal cells. However, recent studies revealed that ASE were abundant such that a huge number of transcript isoforms were found (Kahles et al. 2018). But the functional sequel of most of them were unknown.

Non-directional RNA-sequencing was used in TCGA, so the orientation of all RNA reads was not available. Due to these difficulties, we focused exclusively on the expression level of (directional) junctional reads, which span the exon-exon junctions and directional information was provided by the splicing donor and acceptor sites (GT and AG) of the intron sequence. We profiled the expressed gene junctions of known transcripts and discovered novel junctions in RNA-seq data downloaded from the TCGA database. We handled BAM files from more than 3,000 cancer transcriptomes to identify directional junctional reads to represent various ASEs. There are more than 240,000 novel junctional reads identified. However, the functional sequels of most mRNA isoforms are not known. It could range from neutral due to addition of a few amino acids to a complete change of function if functional domain is lost.

nt-RNA is the only type of mRNA isoform that is certainly non-functional (Figure 1), which could result from various ASE. However, those ASE result in toxic junctions which induce a frame-shift of the ORF of CDS contribute to the majority of nt-RNA transcripts. Our analysis revealed ∼22,000 such toxic junctions, on average 1 per gene. These toxic junctions can be used as signatures to study and quantify the abundance of nt-RNA in various samples.

### 2.2 RNA-seq Data

The RNA-seq data of over 3,000 cancer transcriptome data were obtained from The Cancer Genome Atlas database (TCGA, http://cancergenome.nih.gov/) together with 1,000 normal samples for 13 cancer types. The cancer types include lung adenocarcinoma (LUAD), lung squamous carcinoma (LUSC), breast carcinoma (BRCA), kidney renal clear cell carcinoma (KIRC), kidney renal papillary cell carcinoma (KIRP), kidney chromophobe (KICH), stomach adenocarcinoma (STAD), bladder carcinoma (BLCA), colon adenocarcinoma (COAD), liver hepatocellular carcinoma (LIHC), prostate adenocarcinoma (PRAD) and head and neck squamous cell carcinoma (HNSC). In addition, the RNA-seq datasets of cancer cell lines of 10 cancer types were downloaded from The Cancer Cell Line Encyclopedia (CCLE) project (http://www.broadinstitute.org/ccle/home/). All the RNA-seq data were in BAM file format in which reads have been aligned to the human genome (hg19) by various alignment tools (Table 1).

**Table 1.** The number of the TCGA and CCLE samples and the used RNA-seq aligner used.

| TCGA<br>acro<br>nym | Cancer type | CCLE |  | TCGA normal |  | TCGA tumor |  |
| --- | --- | --- | --- | --- | --- | --- | --- |
|  |  | aligner | No | Aligner | No. | Aligner | No. |
| **BLCA** | Bladder Urothelial Carcinoma | Tophat | 23 | MapSplice | 19 | MapSplice | 286 |
| **BRCA** | Breast Invasive Carcinoma | Tophat | 47 | MapSplice | 110 | MapSplice | 319 |
| **COAD** | Colon adenocarcinoma | Tophat | 41 | MapSplice | 41 | MapSplice | 293 |
| **HNSC** | Head and neck squamous cell carcinoma | Tophat | 27 | MapSplice | 43 | MapSplice | 424 |
| **KIRC** | Kidney renal cell carcinoma | Tophat | 22 | MapSplice | 127 | MapSplice | 425 |
| **LIHC** | Liver hepatocellular carcinoma | Tophat | 30 | MapSplice | 50 | MapSplice | 187 |
| **PRAD** | Prostate adenocarcinoma | Tophat | 4 | MapSplice | 50 | MapSplice | 277 |
| **STAD** | Stomach adenocarcinoma | Tophat | 23 | BWA | 33 | BWA | 83 |
| **THCA** | Thyroid carcinoma | Tophat | 11 | MapSplice | 59 | MapSplice | 268 |
| **LUSC** | Lung squamous cell carcinoma | Tophat | 149 | MapSplice | 107 | MapSplice | 213 |
| **LUAD** | Lung adenocarcinoma | NA | NA | MapSplice | 107 | MapSplice | 303 |
| **KICH** | Kidney Chromophobe | NA | NA | MapSplice | 127 | MapSplice | 66 |
| **KIRP** | Kidney papillary carcinoma | NA | NA | MapSplice | 127 | MapSplice | 151 |

### 2.3 Quality Control workflow

#### 2.3.1 Sample Quality Control

(i) Initial round

BAM files of some samples contained reads of low quality which may be due to sample degradation or laboratory procedures. A stringent sample QC was carried out and samples of good QC were selected as the discovery dataset for datamining for novel splicing junctions. First, samtools (Li et al. 2009) was used to exclude (1) reads unmapped to the genome (script: "samtools -F4"), (2) reads mapped to multiple locations ("samtools -F 0x100"), (3) reads with map quality lower than 40 ("samtools -q 40") and (4) duplicated reads due to PCR bias ("samtools -rmdup"). After removing low-quality reads and duplicated reads, samples with remaining read-depth lower than 40M were also excluded.

(ii) Division into Discovery (higher quality) and Verification sample subsets Samples that passed such initial QC requirements were then divided into (A) discovery datasets for in-depth datamining and quantification of novel splicing junctions and (B) verification datasets, which were used only for quantification of splicing junctions (both known or novel found in the discovery datasets).

Samples were included into the discovery subset if they met all the following criteria:

1. Read depth > 80 M after exclusion of reads by the initial QC
2. Duplication reads ratio was less than 0.4 (40%)
3. Per sequence quality module passed using the Fastqc software
4. Samples should be within the expected range (to exclude outliner samples) of the following 3 criteria:
5. Within the mean ± 2 standard deviation (SD, μ±2SD) of the % of known junctions were expressed, according to normal/cancer tissue type
6. Within the mean ± 2 standard deviation (SD, μ±2SD) of the % of reads mapped to known junctions, according to normal/cancer tissue type
7. Within the mean ± 2 standard deviation (SD, μ±2SD) of the insert size of the paired end reads, according to normal/cancer tissue type Remaining samples not qualified to the discovery dataset were included in the verification dataset. The number of the samples in discovery datasets and verification datasets are shown in Table 2.

**Table 2.**
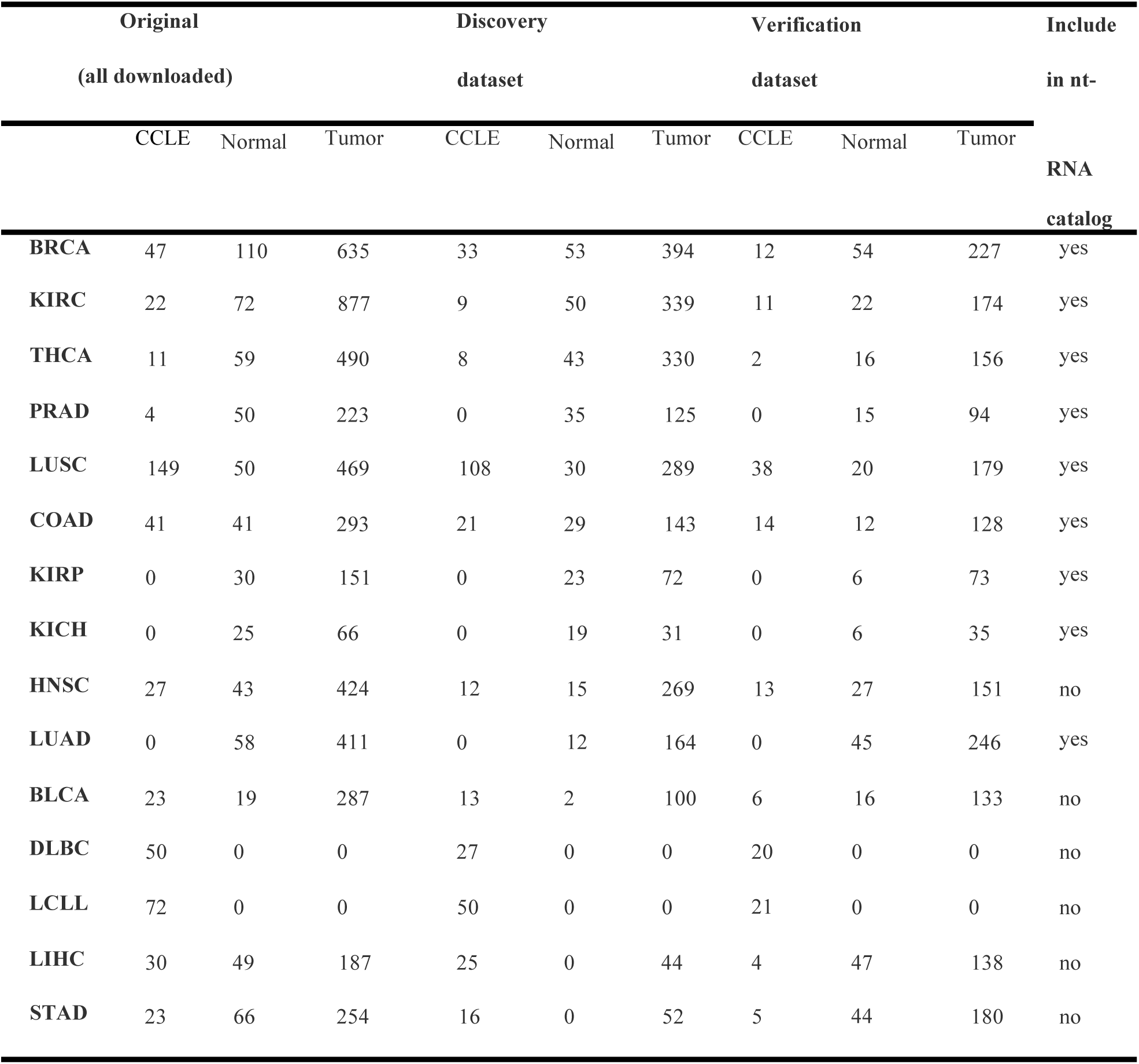
The number of the TCGA and CCLE samples and the RNA-seq alignment algorithms used by TCGA. (e.g. BRCA tumor data of 635 samples were downloaded, in which 394 fulfilled criteria for discovery dataset and 227 fulfilled criteria for verification dataset and 14 samples are not used due to poor QC.)

#### 2.3.2 Sequencing Reads (junctional reads) Quality Control

RNA sequencing of TCGA were performed on the un-stranded library, that is, there was no strand information, and the library preparation was not directional. Due to the loss of the strand information, it is hard to know the strand information of the reads, which may lead to the incorrect quantity of the anti-sense genes whose location have overlap with another protein-coding gene on the opposite strand. Fortunately, we could infer the strand of the junction reads based on the sequence nearing the splicing sites. Thus, in this study, we mainly focus on the expression level of exon-exon junctions reads.

i. Minimal standard

For the determination of steady-state abundance (expression level) of junctional reads in samples, we only consider reads that met all the following three QC criteria:

1. The minimum flanking region on one side of a junction was 4 bp
2. The average quality score of the two bases of both exons adjacent to the splicing site is larger than 30
3. (ii) Junction reads should be mapped at correct orientation (flag=99,163,147 and 73) Additional QC for sequencing reads used in data mining for toxic junctions of novel nt- RNA

Many novel splicing junctions were found which have not been annotated in the NCBI, Ensembl or Non-coding database, but additional QC were needed to make sure that they were not experimental artefacts.

1. The novel junction must be supported by at least ten high-quality reads in at least one cancer dataset to allow cancer specific nt-RNA to be found. These high-quality reads have to fulfil all the following criteria :
2. The minimum flanking region of the junction reads was 16 bp, therefore, the novel junctions were all supported by at least 32 bp across.
3. They had mate reads with map quality score larger than 40 in the paired-end RNA-seq data to confirm genome coordinate.
4. The insert size of the paired junction reads should be within two standard deviations of the mean (μ±2σ) of all paired reads in the RNA-seq data to exclude outliner reads.
5. To rule out the novel junctions were originated from intron sequences, gene fusion or alignment artefacts,
6. To exclude intron sequence or gene fusion, the novel junctions with 15bp flanking sequence were aligned to the genome and transcript sequence and the novel junctions with more than 90% similarity with the known genome or transcript sequence were excluded (ambiguous genome coordinates of the exon partners).
7. To exclude those novel signature junctions with the sum of the distance of the 5’ splicing site and 3’ splice sites to the corresponding protein-coding isoform junctions equal to zero (alignment position shift).

With such stringent criteria, we discovered more than 240,000 novel junctions not annotated in current genome databases. Some of them were reported recently by Kahles et al (Kahles et al. 2018) using a graph approach to resolve the expression of various transcript isoforms. However, the non-directional nature of TCGA data also led to high proportion of false positive findings. On the other hand, our approach identified unambiguous novel splicing junctions though we did not have information on the rest of the transcript.

As the functional sequels of the majority of ASE are uncertain, we confined to analysis of nt- RNA and their associated toxic junctions which could be used for event quantification and differential analysis between cancers and tissues.

### 2.4 Building a dictionary / catalogue of known and novel nt-RNA and their associated toxic junctions

First, we retrieved all known splicing junctions of both “normal” canonical transcripts and isoforms including known nt-RNA isoforms. From these reported nt-RNA isoforms, toxic junctions which lead to ORF frame-shift were identified and entered into a dictionary/catalog of known toxic junctions which will be used for quantification and counting in each samples. We compared the nt-RNA with all the NM transcripts and obtained the positions of the unique (signature) junctions for each nt-RNAs. Afterwards, we also identified the new (novel) toxic junctions in the discovery dataset of each tumor.

#### 2.4.1 Known nt-RNA and their toxic junctions

The annotated transcriptome sequence and transcript isoforms were first retrieved from both the NCBI, Ensembl and NONCODE genome databases. The gene annotation files are downloaded from NCBI (ftp://ftp.ncbi.nlm.nih.gov/genomes/H_sapiens/ARCHIVE/ANNOTATION_RELEASE.105/GFF/ref_GRCh37.p13_top_level.gff3.gz) and Ensembl (ftp://ftp.ensembl.org/pub/grch37/release-84/gtf/homo_sapiens/Homo_sapiens.GRCh37.82.gtf.gz) database (hg19). The noncoding annotation file (hg38) was downloaded from the NONCODE database (http://www.noncode.org/datadownload/NONCODE2016_hg38.lncAndGene.bed.gz) and was remapped to hg19 using liftover software. All the transcripts with CCDS ids were downloaded from the Ensembl database.

Transcript isoforms of a protein-coding gene are defined as the nt-RNA if they fulfil any one of the following criteria:

1. (non-protein coding designation in Ensembl) The transcript has the same frame as the canonical protein-coding gene, but the transcript itself is not labeled as "protein coding" in Ensembl
2. (truncated protein in NCBI) The protein-coding transcripts don’t have CCDS ids, and the length is less than 75% of the longest NM isoform in NCBI
3. (NR_ accession number in NCBI) The id of the transcript in NCBI with prefix "NR_"

#### 2.4.2 Definition of novel toxic junctions as signatures of novel nt-RNA transcripts

To call a novel junction as a toxic junction leading to frameshift of the ORF of a protein coding gene, a new algorithm to calculate the shifted distance of the novel junction compared to the nearest protein-coding isoforms was developed. It uses bedtools (Quinlan and Hall 2010) to identify all the protein-coding exons that overlap with the reads supporting the novel junctions. We only focus on those novel junctions whose flanking sequence on both sides overlap with the exons in a protein-coding gene which will assign the frame of its ORF. With the frame reference of the known protein gene available, the two ends of a novel junction can be mapped to the frame of ORF and distance (gap) between the 2 ends is calculated as its shifted distance. Also, the location of the novel signature junction related to the CDS region of the protein-coding isoforms was also calculated. If the novel signature junction is located within the CDS region and the shifted distance is not a multiple of 3, this novel signature junction causes a frame shift. It is defined as a toxic junction and transcripts carrying this toxic junction are nt- RNA which cannot produce a protein and may be targeted for NMD depending on the location of the toxic junction. A collection of ∼22,000 toxic junctions are available in the GitHub.

Abundance quantification of toxic junctions included showing the summary statistics of the raw counts of the toxic junctions (the 50th, 75th, 90th, 95th and max value of each toxic junction) in the dataset. The data have been uploaded in a BED format to the UCSC genome browser (https://genome.ucsc.edu/cgi-bin/hgTracks) for better search function and visualisation. Both known and novel nt-RNA signature junctions are included in the catalogue and are shown separately in the UCSC database.

### 2.5 Abundance quantification and differential analysis using the catalogue of toxic junctions

With the catalogue of known and novel toxic junctions, abundance quantification was carried out of each junction in samples of the discovery and verification datasets. They were expressed as raw count and normalised to the overall gene expression level. A ratio of usage of the toxic junction was calculated using Splice-in Ratio (SiR) which is similar to splicing ratio but in percentage terms. Another quantification indicator of ASE is percentage splice in (pSI), however, it requires counting reads mapped inside exons for calculation. In the context of a non-directional sequencing method, it is not certain whether inside-exon reads are transcribed by the gene of interest or another transcription motif located on the anti-sense genomic DNA.

Differential analysis of the abundance of toxic junctions were compared between normal and cancer tissue. It was also analysed using renal cell carcinoma dataset as examples and how principal component analysis (PCA) of the abundance of toxic junctions could reveal a subset of patients with a different prognosis.

## 3 Results and Discussion

### 3.1 General trend of expression of toxic junctions (and their associated nt-RNA)

#### (A) The proportional expression of toxic junctions

Similar to few other recent publications, we found the expression of non-translated transcripts (nt-RNA) and the associated toxic junctions were very common in the cancer transcriptome. But they were also found in the normal tissue. And they were produced by most of the expressed protein-coding genes based on the RNA-seq data in discovery datasets.

We analysed for the critical factors affecting the abundance of these non-functional transcripts which co-existed with the functional or canonical transcripts. For example, the toxic junction of nt-RNA of voltage-dependent anion-selective channel 1 (*VDAC1*) which plays an essential role in cancer cell differentiation (Arif et al. 2016) was highly expressed in breast cancer (BRCA samples) and breast cancer cell lines (CCLE samples). Then a linear correlation was found between counts of the toxic junction and total counts of all exon junctions of this gene (Figure 2). The slope in the linear regression plot showed similar slopes for normal, BRCA cancer tissue and CCLE cell lines (Figure 2B). Therefore, the proportional expression of the toxic junction to all junctions of the gene was stable across these 3 types of tissue and it was not affected during oncogenesis. As shown in the Figure 2, one toxic junction transcript was produced for every 70 junction reads in this gene, which gave a ratio of ∼0.014 (i.e. 1.4% of all junction read was a toxic junction). This proportional expression value of a toxic junction can be compared between cancer type and normal tissues (Figure 2C).

**Figure 2.**
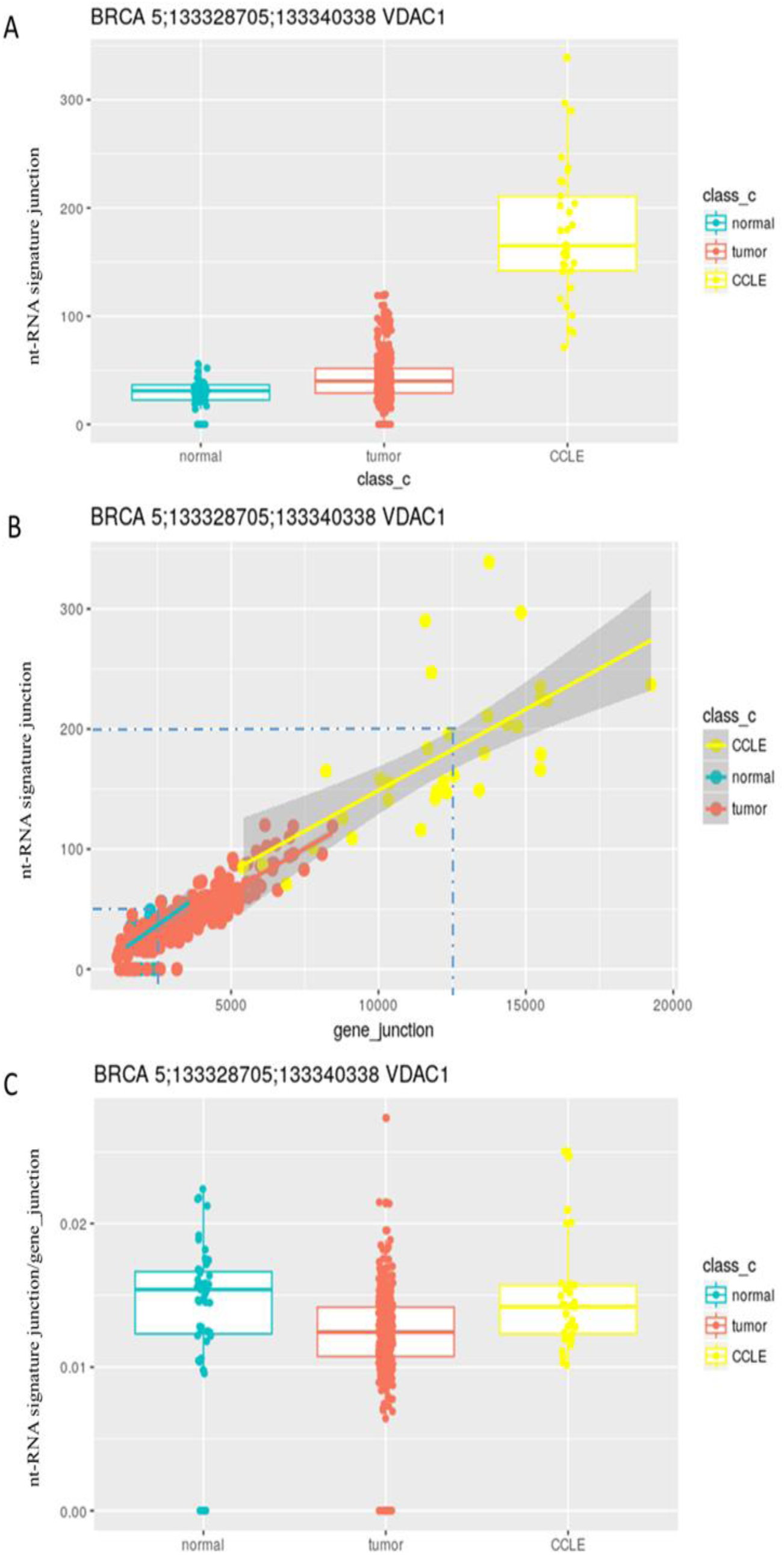
Expression of the toxic junction of gene VDAC1 in Breast normal tissue, tumor **and CCLE samples.** (A) Boxplot of the raw count of the toxic junction. (B) Scatter plot of the toxic junction vs. total VDAC1gene junctions. Although cancer induced (up-regulated) expression of VDAC1, and its nt-RNA, there was no difference in their slopes (proportional expression) (C) Boxplot of the proportional expression ratio of the toxic junctions to the total VDAC1 gene junctions showed the same finding that there was no major difference in proportional expression of this toxic junction of VDAC1 between normal tissue and cancer.

We then used the linear model to investigate the proportional expression of the toxic junctions to all gene junctions of various genes in various cancer and normal tissues. The coefficient (slope) of the gene total junction count represented the average proportional expression of the toxic junction in various cancer types. The distribution of the proportional expression of toxic junctions in various cancer datasets (including normal samples and tumor samples) shows that the highest peak (mode value) was located at around 0.1 for most cancers (Figure 3). It indicates that about 10% of transcripts contained a toxic junction, giving rise to nt-RNA. Interestingly, we have also found that some toxic junctions (thus the associated nt-RNA) in some gene were expressed in large quantities, more than 10% of the transcripts (as shown by the long tail in the distribution in Figure 3). Such a heavily right-skewed distribution was found across all cancer types, and some toxic junctions accounted for more than 50% of all transcripts. The ntRNA in these genes may play a potential role during the tumor progression as half of the transcripts are non-functional, which is reminiscent to the situation of a heterozygous somatic mutation.

**Figure 3.**
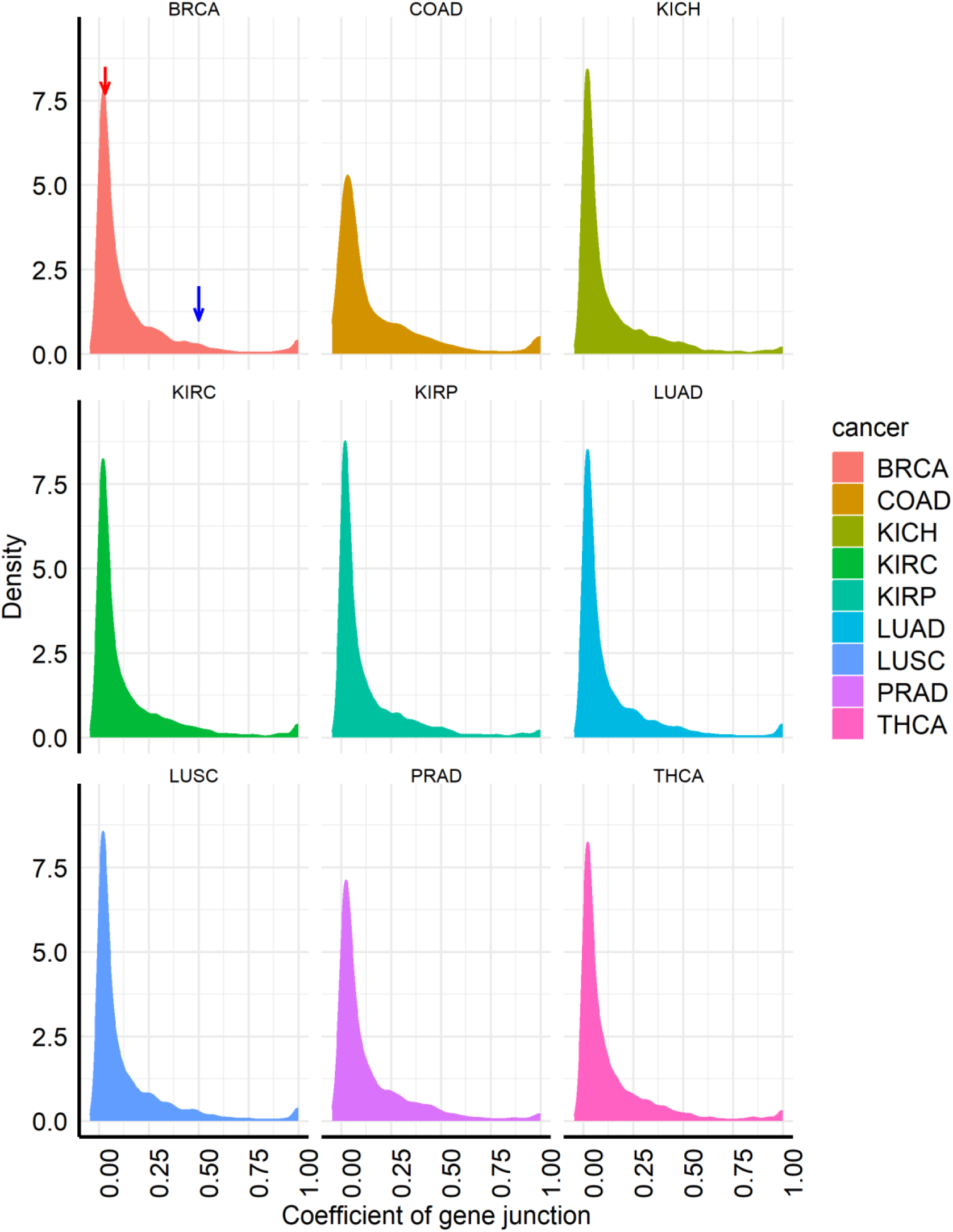
The distribution of the coefficient (slope, representing the proportional expression of nt-RNA) of gene junctions in the linear model of all toxic junction in different cancer datasets (including normal and tumor samples). For example, in BRCA, the highest peak (red arrow) located below 0.1 indicating that most nt-RNA had a proportional expression of around 10%. However, some uncommon genes express nt-RNA as much as 50% of all transcripts (0.5, marked by blue arrow)

#### (B) Expression pattern of the nt-RNAs in cancer and normal samples

For the reason mentioned above, we wanted to know if the proportional expression of toxic junction (nt-RNA) was different between normal and cancer samples. We firstly analyzed the scatter plots of the raw count of the toxic junction and total gene junction and found that the expression of the majority of toxic junctions were correlated with expression levels of the corresponding gene.

Generally, expression pattern of toxic junctions could mainly be grouped into the following 4 categories (Figure 4):

1. Both normal and tumor samples express the nt-RNA. The raw count of the signature junction is highly correlated with the total gene junction counts, and the proportion of expression is similar in both normal and tumor samples. That is slopes were similar.
2. Only normal or tumor samples express the nt-RNA, and its expression ratio is similar among samples.
3. Only a small part of normal or tumor samples express the nt-RNA, and the ratio of its expression is dynamic.
4. Both normal and tumor samples express the nt-RNA, but have different proportional expression of the nt-RNA between cancer and normal tissue. Slopes of normal and tumor samples are different.

**Figure 4.**
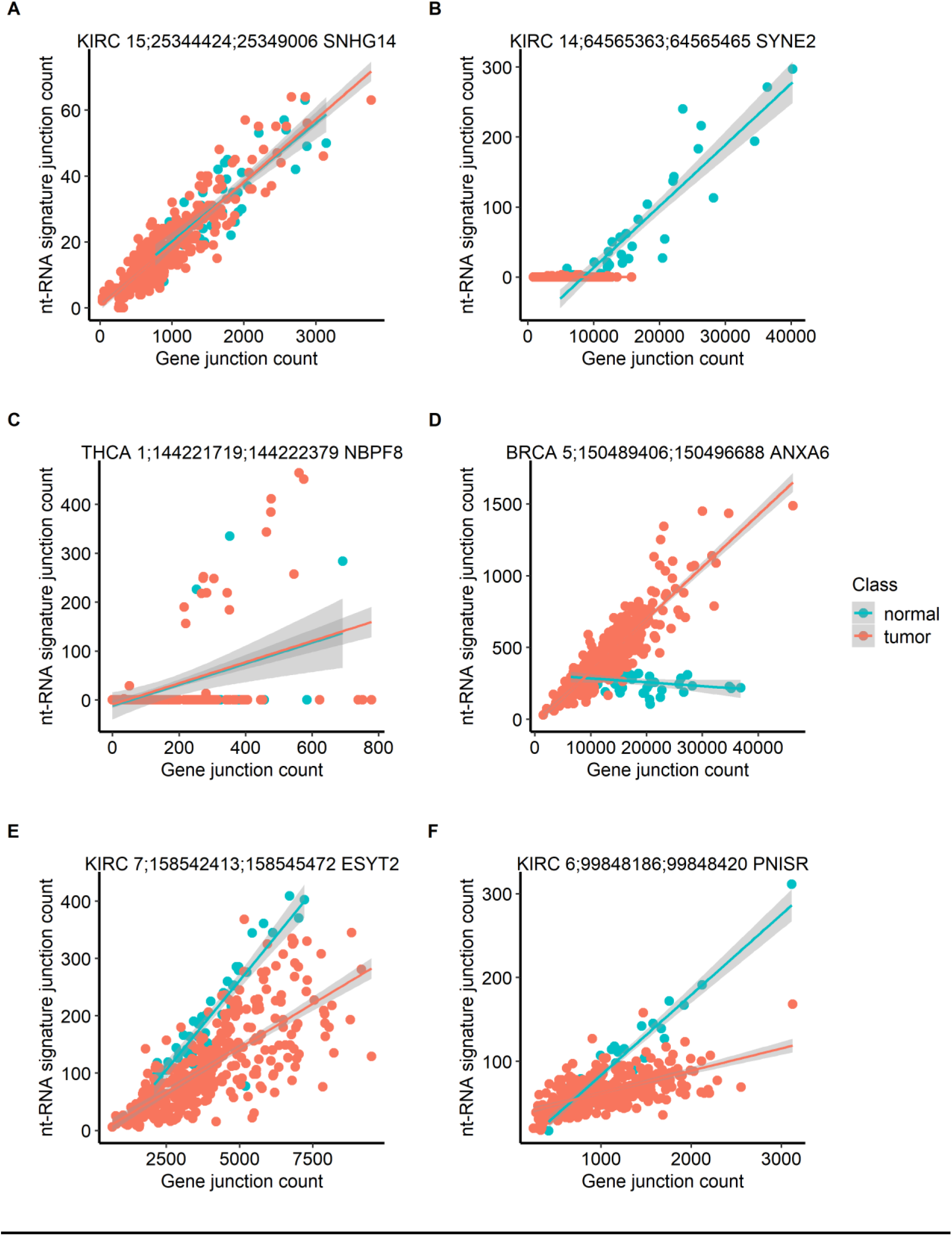
Examples of various patterns of proportional expression of nt-RNAs in normal and tumor samples as shown in scatter plots. The x-axis is the total count of the gene junction and the y-axis is the raw count of the nt-RNA signature junction. Figure 4A is toxic junction in *SNHG1* gene in KIRC as an example of type 1, Figure 4B is *SYNE2* gene in KIRC as an example of type 2, Figure 4C is *NBPF8* gene as an example of type 3 and Figure 4D to 4F are *ANXA6, ESYT2* and *PNISR*, respectively as examples of type 4.

To comprehensively analyze the difference between healthy and cancer samples, in the proportional expression of toxic junction relative to the corresponding protein-coding genes, we used the coefficients of the intercept-terms in the regression model with an interaction term of the gene junction expression and a dummy variable (Class: i.e., normal or tumor) after log transformation of the data to evaluate the difference in the proportional expression of the nt- RNA between normal and cancer samples.

As shown in Figure 5, the closer the intercept coefficient of the cross-terms to 0, there was no difference in proportional expression of the nt-RNA between normal and cancer samples. The intercept coefficient of the most toxic junction in the linear model with interaction term is close to 0 which indicate that the ratio of expression of the most toxic junction is not significantly different between cancer and normal samples. There is a small proportion of toxic junction with different expression ratio between normal and tumor samples, and this difference may be partly because that these nt-RNA genes are only expressed in cancer or normal samples. For those toxic junctions that are both expressed in normal and tumor samples but with different expression ratio, they may play an important role as potential biomarkers.

**Figure 5.**
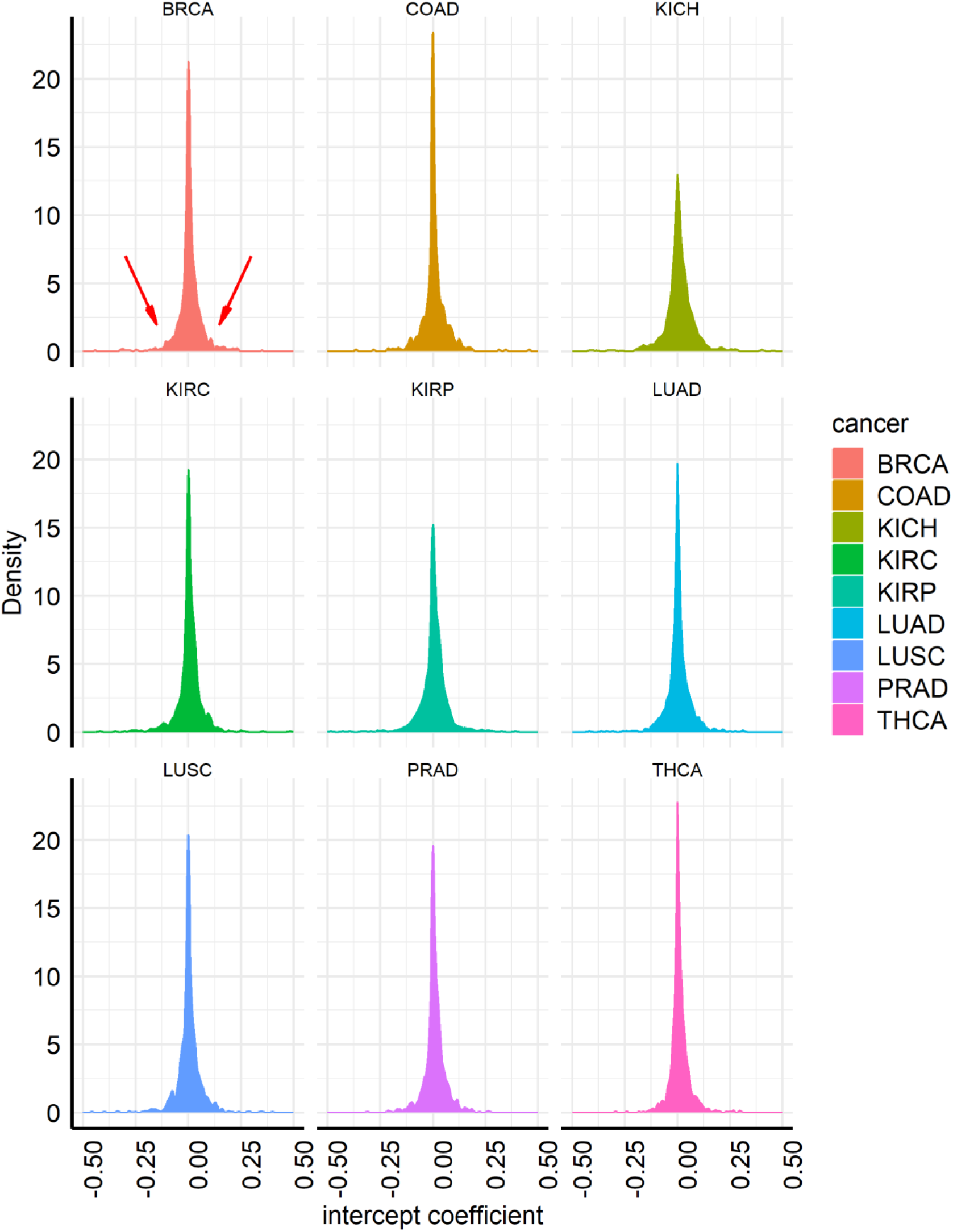
The distribution of the intercept coefficient in the linear model with interaction item of all nt-RNA signature junction in different cancer datasets (including normal and tumor samples). Since the data are log transformed, the intercept coefficient represented the difference of proportional expression of nt-RNA between cancer and normal tissue. The peak (e.g. majority of genes) had intercept coefficient of zero, that is, there was no difference between cancer and normal. However, there were shoulders (e.g. arrows in BRCA chart) on both sides of the peak. There were many genes with differential proportional expression of nt- RNA between cancer and normal.

#### (C) Splicing in Ratio (SiR)

Instead of calculating the slope and using scatter plot to compare the proportional expression of nt-RNA, this proportion expression could be expressed as a single value, Splicing in Ratio (SiR). We divided the raw count of the nt-RNA by the sum of the fragments aligned to its 5’ splice site and the 3’ cleavage site (as shown in Figure 6) to obtain the splicing in ratio (SiR) of each toxic junction. This value allows us to evaluate the nt-RNA expression ratio more accurately. However, there is a limitation, for signature junctions which do not share same 5’ splicing sites or 3’s splice sites with other expressed junctions, their SiR will be 1 and is not meaningful. The expression ratio of these toxic junctions can only be calculated by dividing the raw count by the total gene count. SiR is similar to another index called Splice Ratio with standardisation to 100%. Percentage splice in (pSI) is another commonly used parameter, however, pSI is adversely affected by non-directional RNA-sequencing data in which the strand origin of exon-reads cannot be unambiguously assigned.

**Figure 6.**
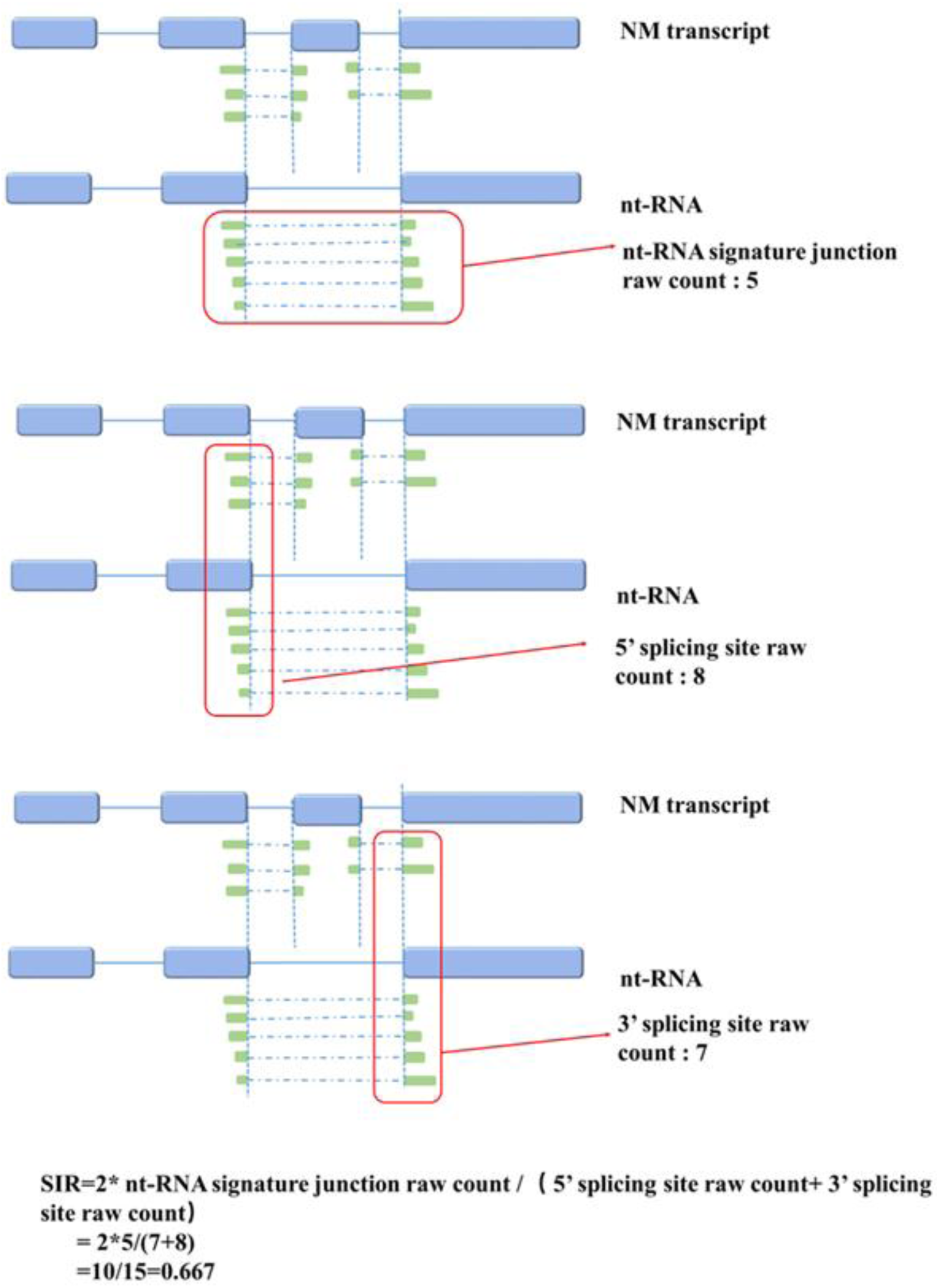
**Calculation of the Splicing in ratio (SiR)**; the light blue box represents exons and lines between the light blue box is the intron. The green box represents spliced reads aligned to the junctions in the RNA-seq data.

### 3.2 Examples of differentially expressed nt-RNAs

#### (A) nt-RNA in ANXA6

Annexin VI (ANXA6) is a member of a highly conserved family of calcium-dependent phospholipid-binding proteins and play a role in calcium homeostasis, membrane traffic, and membrane organization. ANXA6 could act as a regulator of the epidermal growth factor receptor and the Ras signaling pathway(Grewal et al. 2010). Recent publications show that ANXA6 may exhibit different functions in the different stage of Breast cancer and various breast cancer cell lines (Qi et al. 2015). The expression of ANXA6 could increase the invasiveness of the Breast cancer while its downregulation will enhance the growth of some Breast cancer cell lines (de Muga et al. 2009; Sakwe et al. 2011; Koese et al. 2013). This gene will generate two major isoforms (AnxA6-1 and AnxA6-2) due to alternative splicing. The ANXA6-1 was predominately expressed in most tissues. The Anxa6-2 skipped the exon 21 and lost the VAAEIL sequence. The proportional expression of the ANXN6-2 expression is higher in breast cancer than that in normal samples (Wilcoxon text: P-value=1.805e-15). We found that only a small amount of ANXN6-2 is expressed in normal breast samples, and its expression is not correlated with the expression of the entire gene. However, in breast cancer samples, the ANXN6-2 expression is accounted for more than half of the gene expression, and it is significantly positively correlated with the overall expression of the entire gene. It was reported that these two isoforms have independently different function and location. For example, EGF- dependent Ca2+ influx in A431 cells is only inhibited by ANXN6-1 (Fleet et al. 1999), while ANXN6-2 has a stronger affinity for Ca2+ and a stronger F-actin binding (Strzelecka-Kiliszek et al. 2008). However, the functions of these two isoforms during the development of breast cancer have not been further studied in the current studies.

Our results indicate that ANXN6-1 is predominantly expressed in normal breast cells, whereas ANXN6-2 are predominantly expressed in breast cancer samples. This result suggests that ANXN6-2 with different functions are needed in the development of cancer, and it may play an essential role during the development of breast cancer. In-depth study of the isoform switch of ANXN6 between breast normal and tumor samples and its functions may provide a better understanding of the mechanism of breast cancer development.

#### (B) Toxic junctions (and nt-RNAs) in ESYT2

The protein of gene ESYT2 Extended synaptotagmin-2 has been demonstrated to interact with the Fibroblast Growth Factor Receptor and activated FGF receptor (Jean et al. 2010; Tremblay et al. 2015). It plays a vital role in growth factor signaling. This gene mainly expressed two isoforms: a short isoform (ESYT2-S) and a long isoform (ESYT2-L: nt-RNA) which include a cassette exon between exons 13 and 14. We observed a significant isoform switch of ESYT2 based on the RNA-seq data of the renal cell carcinoma (KIRC) samples. The expression ratio of the long ESYT2 isoform (ESYT2-L) to the short isoform (ESYT2-S) is higher in normal kidney samples than the tumor samples (Wilcoxon text: P-value < 2.2e-16). The Kaplan-Meier survival curves showed that samples with the higher expression ratio of ESYT2-L are associated with better survival (p=2.04e-06). Multivariate Cox proportional hazards regression revealed that the expression ratio of the ESYT2-L could be as an independent prognostic factor for patients with CRC (hazard ratio, 0.037; 95% confidence interval, 0.01-0.125; P=1.24e-07). Besides, the Gene set enrichment analysis (GSEA) indicated that genes correlated with the expression ratio of ESYT2-L are enriched in the hallmark of the EMT and invasiveness signature from cancer cell. Our findings show that the alternative splicing of ESYT2 could be a potential prognostic biomarker in KIRC and samples with a lower expression ratio of the ESYT2-L isoform may be more likely to have the potential to become metastatic.

#### (C) Toxic junction in ERBB2 affects prognosis

We also calculated the SiR for the novel junction and found some novel junctions that are highly expressed in tumor samples compared with the normal samples. For examples, ERBB2 encodes a member of the epidermal growth factor (EGF) receptor family of receptor tyrosine kinases. It is a well-known oncogene. We found that novel isoform, which has a novel junction with an alternative 3’ splicing site in exon 16, is highly expressed in BRCA tumor samples. The Kaplan-Meier survival curves showed that samples with higher SiR of the new isoform in ERBB2 are associated with better survival (p-value=0.0153). This result suggests that in addition to nt-RNA, new isoform also could be used as a potential biomarker.

#### (D) Toxic junction of TP53 was related to somatic mutation in the gene

Multiple splicing factors are involved in RNA splicing. Abnormal expression of these splicing regulatory factors may induce alternative splicing. However, the mechanism by which these proteins regulate splicing is complicated and unclear. Besides the abnormal expression of splicing factors, mutations is another reason that causes alternative splicing. Somatic mutations causing non-sense, frame-shift and premature termination are found in various cancers (Seiler et al. 2018). Also, mutations close to the splice site have a very high probability of causing abnormal splicing (Krawczak et al. 2007). To analyze the effect of the mutations on alternative splicing, we downloaded the somatic mutation datasets from the TCGA datasets.

TP53 is an essential tumor suppressor gene. There are 28 known transcript variants for TP53. Besides, we identified seven novel transcripts. 71% of them led to a frame-shift in the protein coding sequence. The transcript causing a splicing aberration between exon 7 and exon 8 was recurrent in 30 tumor samples. These tumors were originated from breast, lung, colon and thyroid tissues. We then investigated the mechanism responsible for this splicing event. All of the samples with the high expression level of this novel transcript could be explained by one of the two somatic mutations near the splicing sites. Namely, a C to A mutation at the splicing acceptor site and a C to A/G mutation at the 6bp downstream of the end of intron 7. Finally, we compared the survival of patients with or without this splicing aberration. We found that breast cancer patients without splicing aberration have better survival.

In addition to TP53, we also found many samples that have highly expressed toxic junction in gene CDKN2A and HLA-DPB1 tend to have somatic mutations. Our data indicate that somatic mutations may be the reason that leads to alternative splicing events in a few samples. For alternative splicing that recurrent in most samples without somatic mutations in the nearby region, it may be regulated by more complex mechanisms.

In this paper, we mainly studied the efficiency of the generation of the toxic junction and novel signature junction based on their expression profiles of different cancer and tissue samples obtained. We calculated the splicing in ratio (SiR: raw junction count divided by the sum of the reads aligned to 5’ or 3’ splice site) for each junction to estimate how efficiently the junctions will be generated in each sample. And we found some nt-RNA or novel isoform could be used as potential prognostic biomarkers by comparing the SiR values of the signature junctions between normal samples and tumor samples. We also found that somatic mutations may result in abnormally expressed transcripts with high expression value in a small number of samples.

In summary, analysis of RNA-seq of cancer transcriptome provides essential information about the gene expression and variation of transcripts. It also provides a means to assess the functional consequence of somatic mutations and characterize novel transcripts. This approach enables identification of novel cancer biomarkers with nt-RNA junctions and provides a mathematical framework to show the dynamics of production of nt-RNA junctions.

### 3.3 Summary statistics of about 22,000 toxic junctions and novel junctions found by this study embedded in the UCSC Genome browser

More than 250,000 ASE junctions were revealed by our datamining of TCGA pan-cancer transcriptomes. ASE occurred within 1000 bp of a known splicing site where the splicing was supposed to occur (Figure 7A). The classification of ASE junctions is shown in Figure 7B. 49% of ASE junctions do not map to any known junction at either splicing donor or acceptor of known introns (e.g. intron retention).

**Figure 7.**
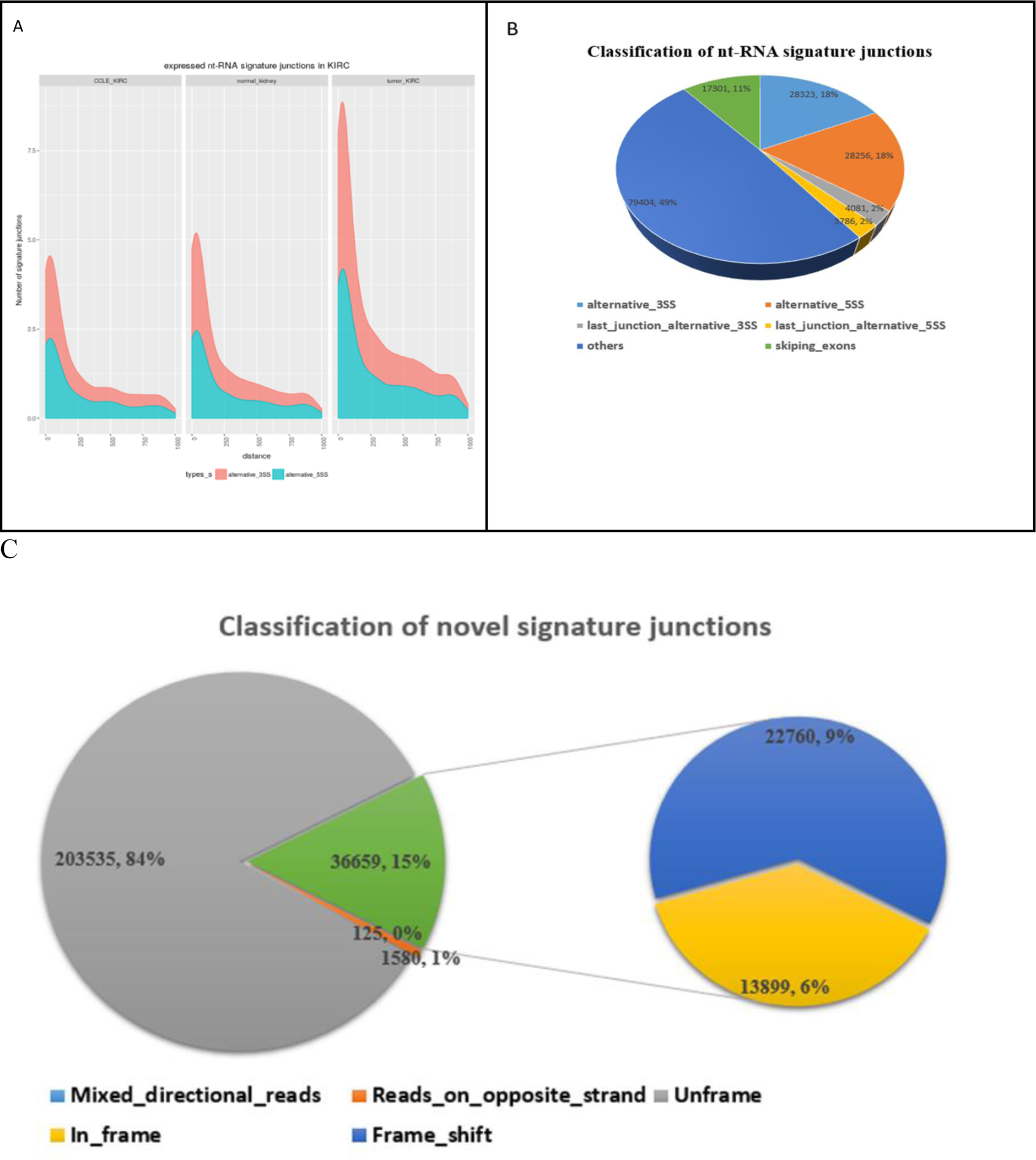
Characteristics of ASE junctions found in pan-cancer transcriptomes**. A.** Density distribution of toxic junctions by distance from the nearest known “normal” NM junction. Most of them are within a distance of 500 bp. Cancer tissue (e.g. KIRC) expressed several times more toxic junctions than normal tissues. B. Distribution of ASE revealed from pan-cancer transcriptomes. C. 60% of novel (unannotated) junctions are toxic junctions leading to ORF frame shift and production of non-translated RNA (nt-RNA).

For all junctions mappable to one known splicing donor or acceptor (51%), they are divided into 3 major groups: 1. alternative 5’ donor site (18%), 2. alternative 3’ acceptor site (18%) and 3. Exon skipping (11%). As expected alternative splicing site in the first exon (2%) and alternative site at the last exon (2%) accounted for a minority of the ASE. 15% of ASE were novel, which were not annotated in Genome databases (Figure 7C). In turn, almost two-thirds (∼60%) of the novel ASE (represented by 22,760 toxic junction) resulted in a frame-shift of ORF. Therefore, unannonated splicing junctions may be an important regulator of biology and oncogenesis. Therefore, we archived their summary statistics into GitHub and their locations are open to researchers to look up in the UCSD Genome Browser (Figure 8).

**Figure 8.**
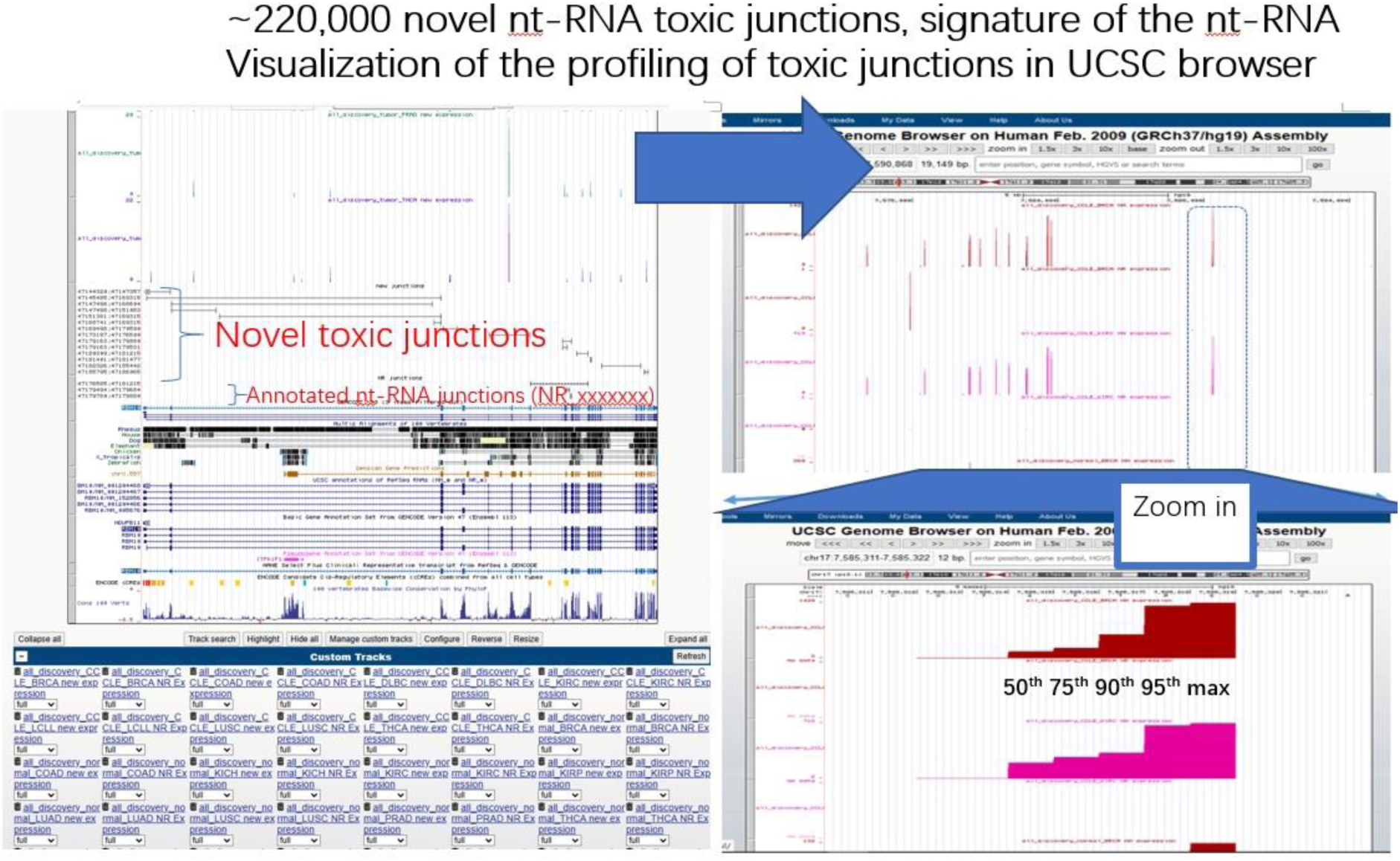
shows the appearance of the archive of ∼22,000 novel toxic junctions in UCSC browser. The novel toxic junctions and known toxic junctions from annotated transcripts in Genome databases (e.g. with NR_xxxxxxx accession in NCBI) are embedded in the Genome Map (Left panel). The selection option of UCSC browser allows users to choose which caner types to show. Summary statistics including raw read counts and SiR are shown as the median, 75th, 90th, 95th percentile and maximal values for each cancer and normal tissue type for each toxic junction

### 3.4 Unsupervised clustering of toxic junctions defined new subtype of RCC in KIRC samples based on the SiR value of highly expressed junctions

We measured the steady-state abundance of ASE by using SiR value of each junction. Besides, by comparing the difference in the SiR value of junctions between normal and tumor samples, we found junctions in gene ESYT2, and ERBB2 might be used as potential biomarkers. We investigated by multivariate statistics to determine whether SiR values of toxic junctions affected cancer survival. The main focus was on the discovery datasets of KIRC tumor samples and normal samples.

Renal Cell Carcinoma (RCC) is a tumor with a high degree of malignancy and a high mortality rate among urinary system tumors. There are three common subtypes of renal cancer based on their histological characteristics: clear cell renal cell carcinomas (ccRCC), also as KIRC in short, papillary renal cell carcinomas (pRCC), and chromophobe renal cell carcinomas (crRCC). Among them, clear cell type renal cell carcinoma is the most common and malignant subtype of renal cell carcinoma, accounting for 70% to 80% of renal cancer (Rini et al. 2009) Because Renal Cell Carcinoma is usually not symptomatic at an early stage, most cancer patients cannot be detected early and cause delays in treatment. Therefore, if we could identify the cancer-related molecular markers in the early stages of tumor formation in which the patient has no clinical symptoms, the probability of the early diagnosis of cancer can be significantly improved. Also, due to individual genetic differences, patients’ responses to drugs are also different, identifying different cancer subtypes by molecular markers can help develop drugs for different subtypes in the clinic, thereby improving the therapeutic effect and prognosis.

We first calculated the raw count and SiR values for all junctions in the KIRC normal and cancer samples. Next, we selected those candidate junctions that were both moderately expressed and showed variable expression among the KIRC tumor samples (Figure 9). In total, we included 173 candidate junctions. Based on these junctions, we used unsupervised clustering methods to cluster all KIRC tumor samples (all belonged to ccRCC) in the discovery datasets. We found that these samples formed four clusters by unsupervised K-means clustering (Figure 10A). Samples in the second and fourth clusters have better survival while samples in the first and third clusters are correlated with worse survival. The purpose of this unsupervised clustering analysis was to see if the collection of these ASE data provided information about cancer survival (Figure 10B). The findings of cluster-related survival confirmed that ASE had important biological significance and were informative for disease prognosis (Figure 10B).

**Figure 9.**
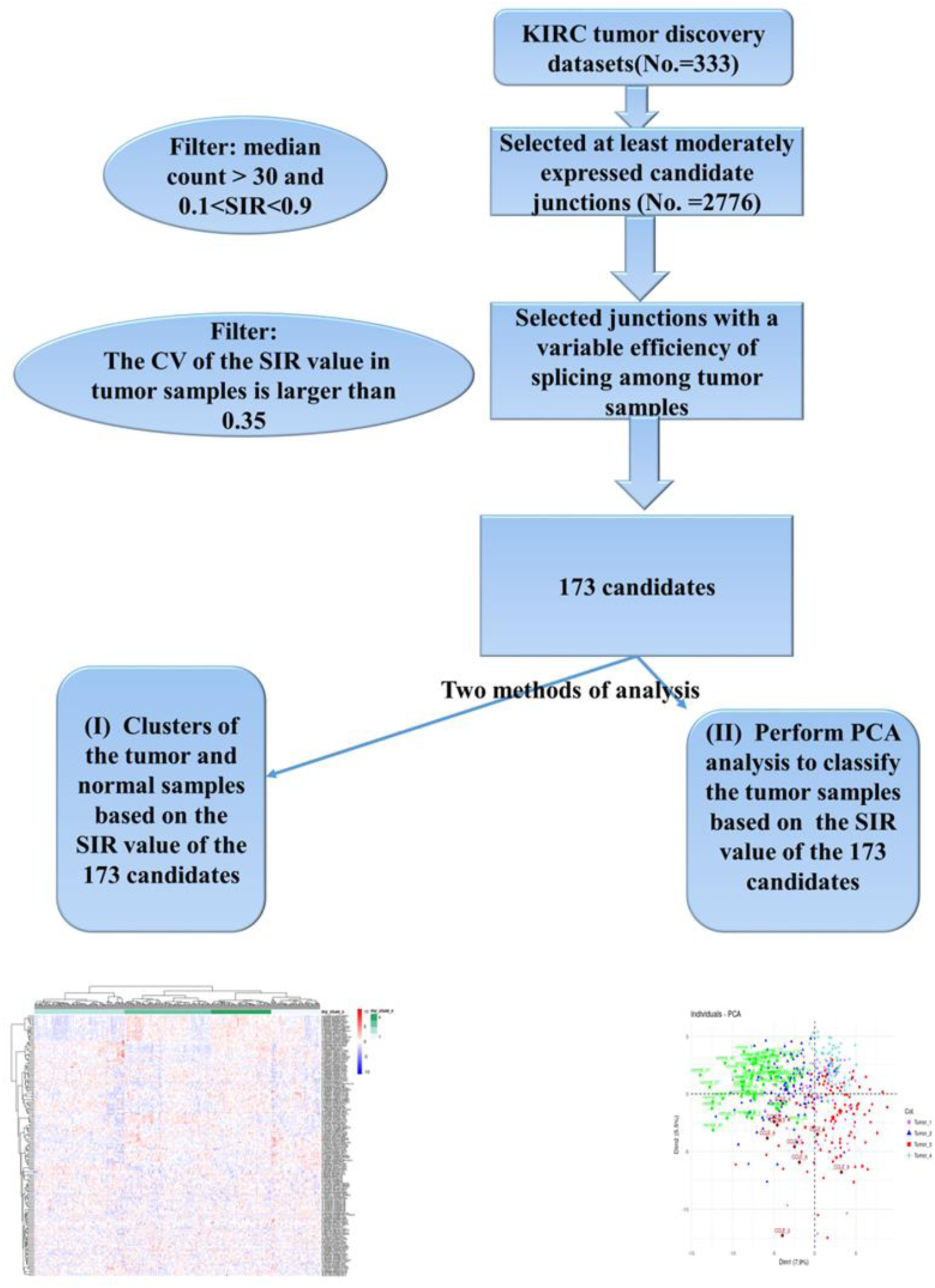
Schematic workflow for analyzing the SiR value of junctions altered in KIRC tumor samples.

**Figure 10.**
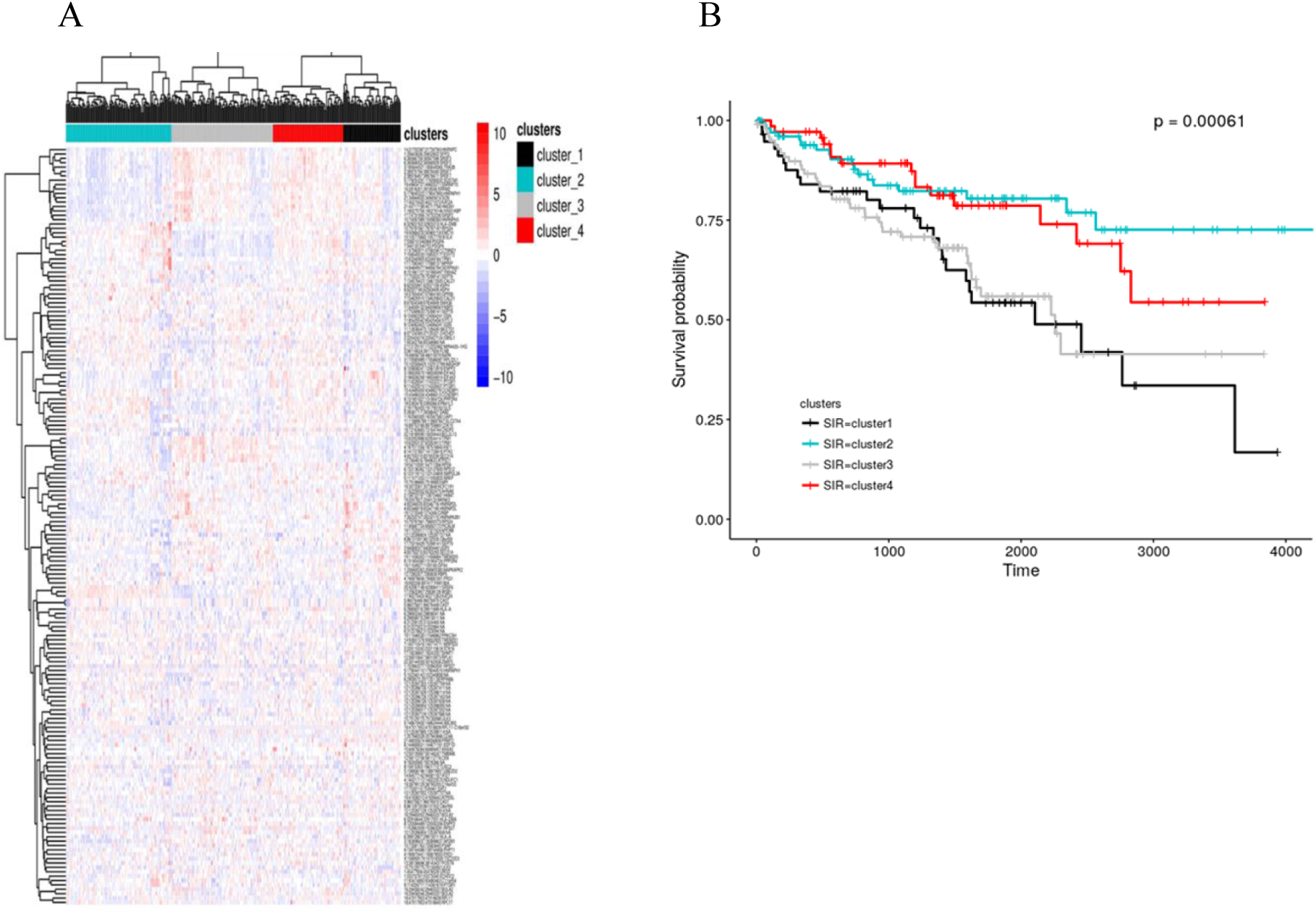
A. Heatmap of the KIRC tumor samples based on the SiR value of the 173 candidate junctions (the name of the genes are show in the row names). B. Survial curve of 4 clusters of samples.

### 3.5 PCA analysis to identify which ASE contribute to prognostic information

The SiR values of these ASE in candidate genes was useful to cluster patients into different prognosis groups. Principal component analysis (PCA) which is a dimensionality reduction method was used to analyse the ASE signals. The top ten principal components (PC) accounted for more than 35% of the total variance. The first PC can account for 7.9% of the variance while the second PC could count 5.4% (Figure 11A).

**Figure 11.**
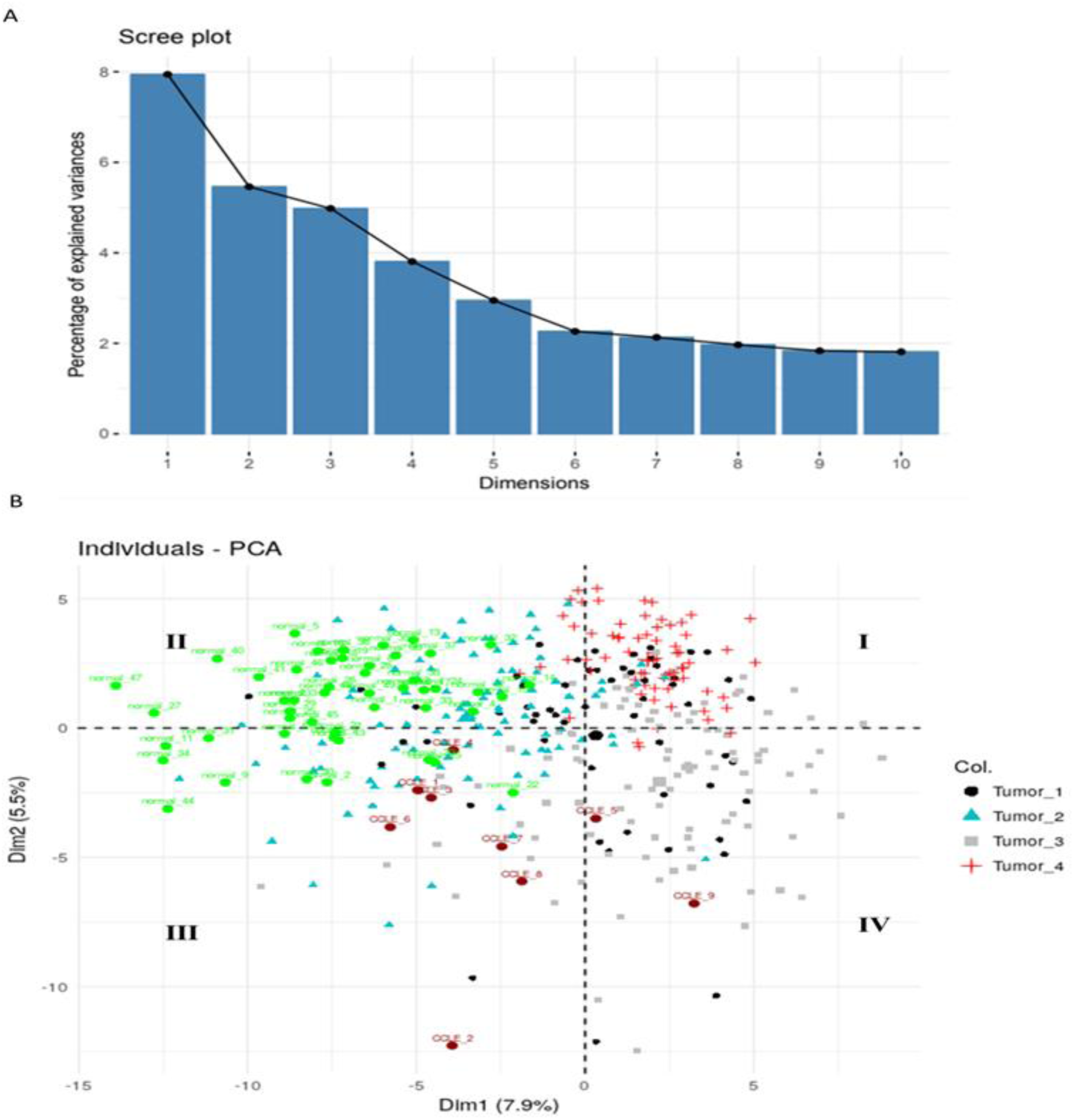
PCA result based on the log-transformed SiR value in KIRC tumor samples. (A) The percentage of explained variance for the top 10 principal components. (B) The biplot of the first two principal components for individuals. The tumor samples are marked by different color and shape based on the clusters. The location of the normal samples (green) and CCLE samples (dark red) are predicted based on the PCA model of tumor samples.

The results of PCA were coherent to that in the unsupervised clustering. The 4 clusters of KIRC samples by unsupervised clustering were plotted with different symbols against the value of PC1 and PC2 (Figure 11B). The samples in the cluster-3 are mainly in the fourth quadrant, while the samples of prolonged survivors in clusters 2 and 4 are primarily in the second quadrant and first quadrant, respectively. Normal samples mostly located in quadrant II as well, together with most of the Tumor_2 cancer. It also coincided that Tumor_2 had the best prognosis. Thus, we performed a survival analysis based on the Cox proportional hazards model on these ten PCs. Under the control of age, gender, grade, VHL mutation and other factors, we found that PC1 and PC2 can be used as an independent diagnostic biomarker. Samples with a higher PC1 value have a worse prognosis. This observation is consistent with the results obtained from clustering. Through the PCA model generated based on cancer, the normal samples were mainly distributed in the second quadrants of the biplot, while CCLE samples were in the third and fourth quadrants. This result indicates that the cancer samples in the second quadrants have a better the prognosis, which may be partly explained by the observation that their ASE properties are similar to that of normal samples. Overall, the PCA results further confirmed the clustering results of junction-based SiR.

Since PC1 and PC2 are associated with prognosis, and they can be used as independent diagnostic biomarkers (Figure 12), we wanted to know which genes have the most significant contribution to PC1 and PC2. We then performed gene enrichment for those top component genes based on the gene sets in the MSigDB database (Liberzon et al. 2011). The top component genes for PC1 are enriched in the gene sets related to RNA splicing (Table 3), while the top component genes for PC2 are enriched with the gene sets which are correlated with the immune cell (Figure 13).

**Figure 12.**
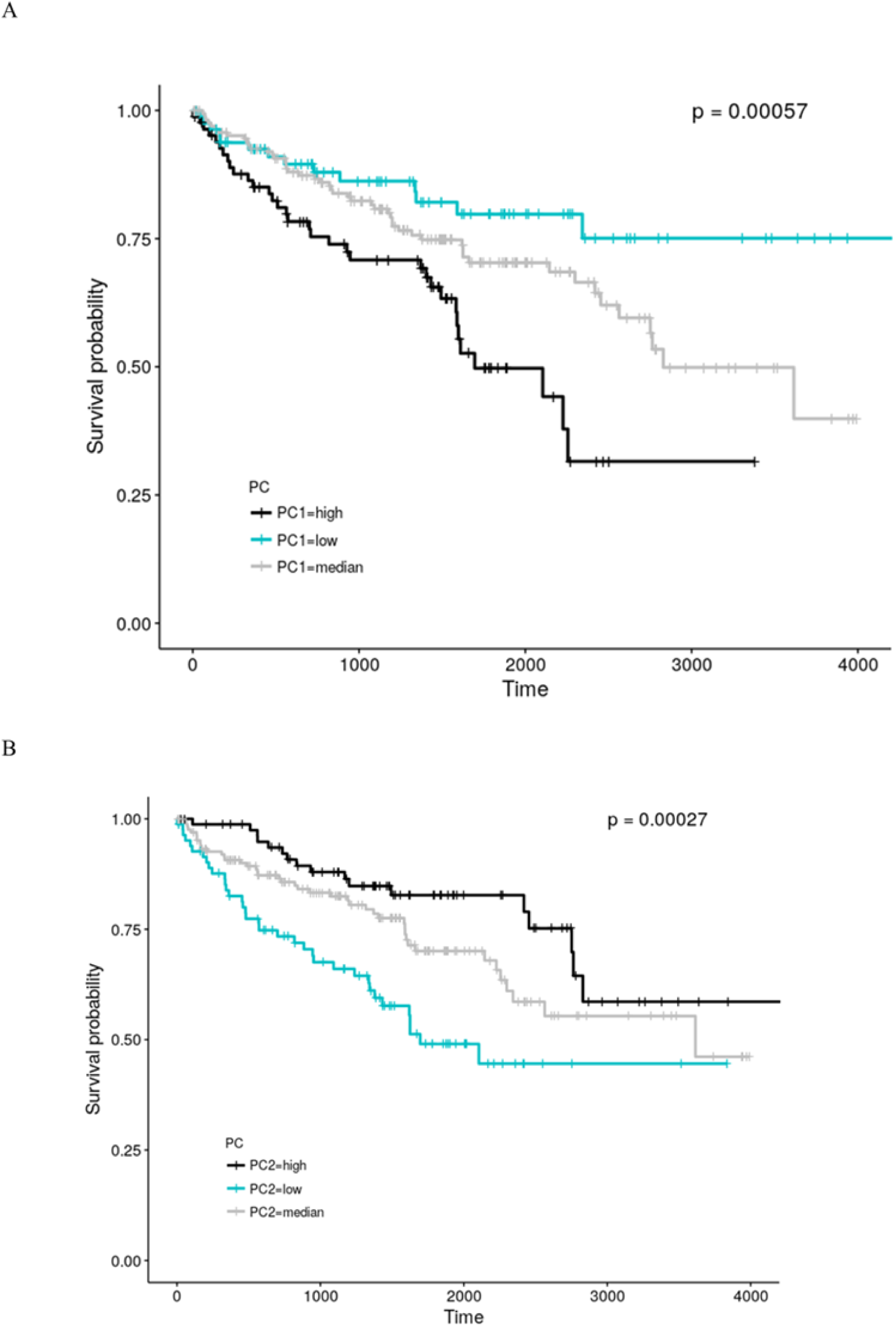
Kaplan-Meier plot for scores of PC1 and PC2 in KIRC tumor samples. High PC1 score and low PC2 score were associated with poor survival.

**Figure 13.**
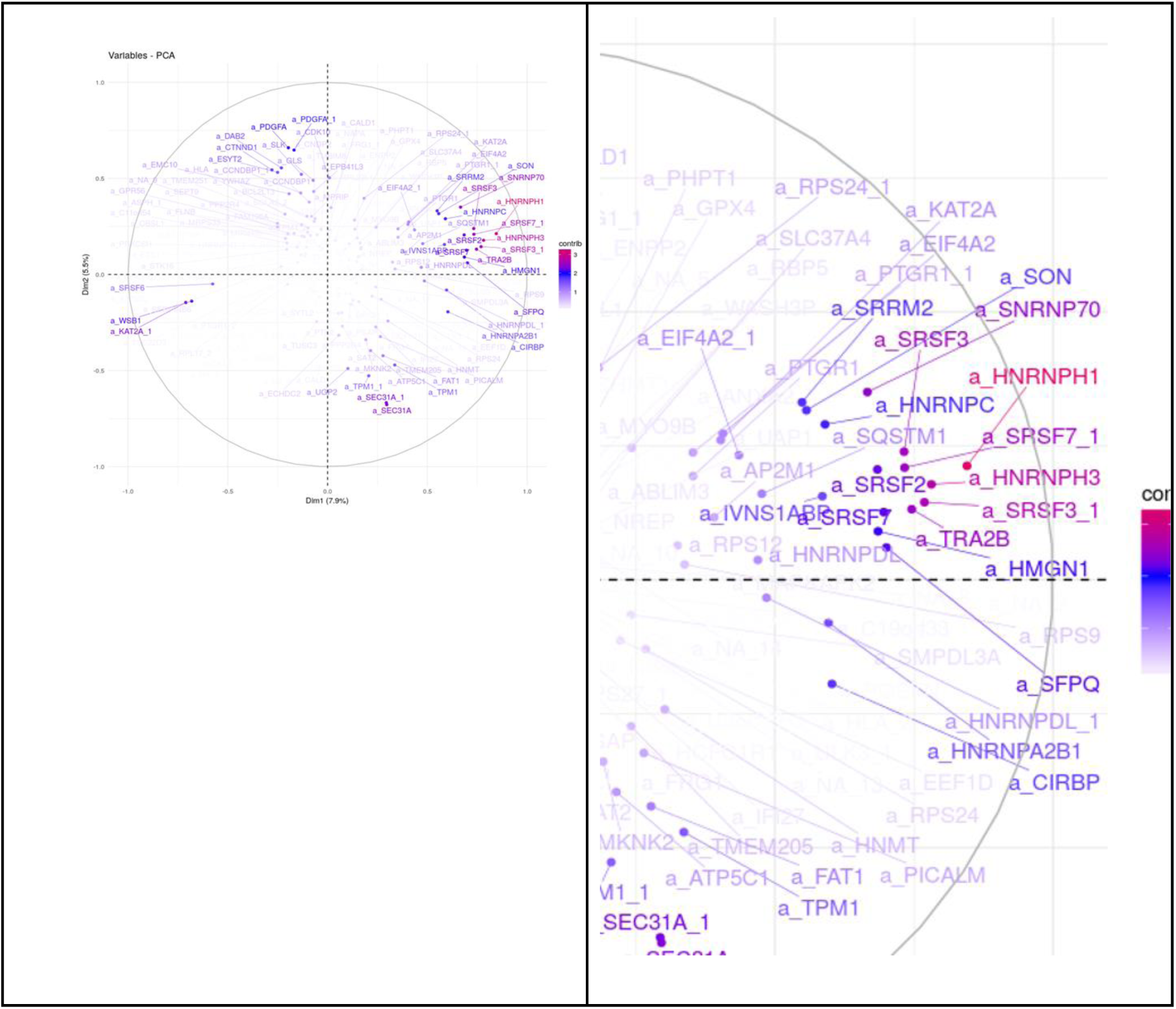
The biplot of the first two principal components of PCA analysis. The contributing component gene in each PCs are shown. For example, PC1 (Dim1, x-axis) has many splicing factors as its components, e.g. SRSF3, SRSF7, and HNRNPH1. They are positively correlated with PC1.

**Table 3.**
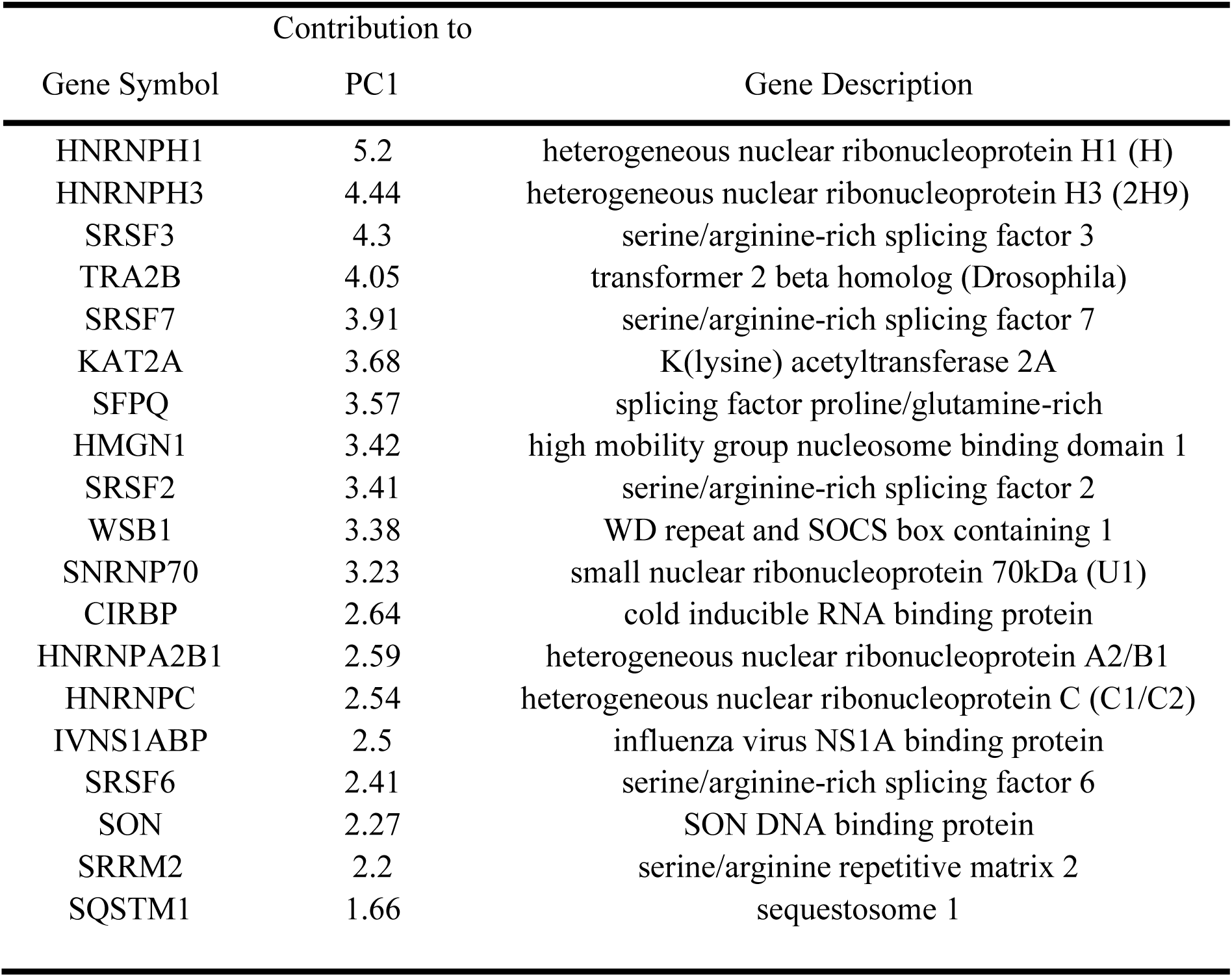
The top component genes on PC1.

#### (A) Principal components and Tumor-infiltrating lymphocytes

Tumor-infiltrating lymphocytes (TIL) have been found in many cancers (Li et al. 2016; Varn et al. 2018), these invading immune cells may have different alternative splicing events or different splicing efficiencies when compared to that in tumor cells. At the same time, the number of invading immune cells may also be related to survival. We then investigated whether PC1 and PC2 are related to TIL. We obtained the data of TIL from these published papers (Li et al. 2016; Danaher et al. 2017). From the scatter plot in Figure 14 we can see, PC1 and PC2 are correlated with the score of some immune cells. For example, PC1 is positively correlated with the score of Type 1 helper T (Th1_cell) (R²=0.41), while the PC2 and score of Dendritic cells (DC) is negatively correlated (R²= -0.58).

**Figure 14.**
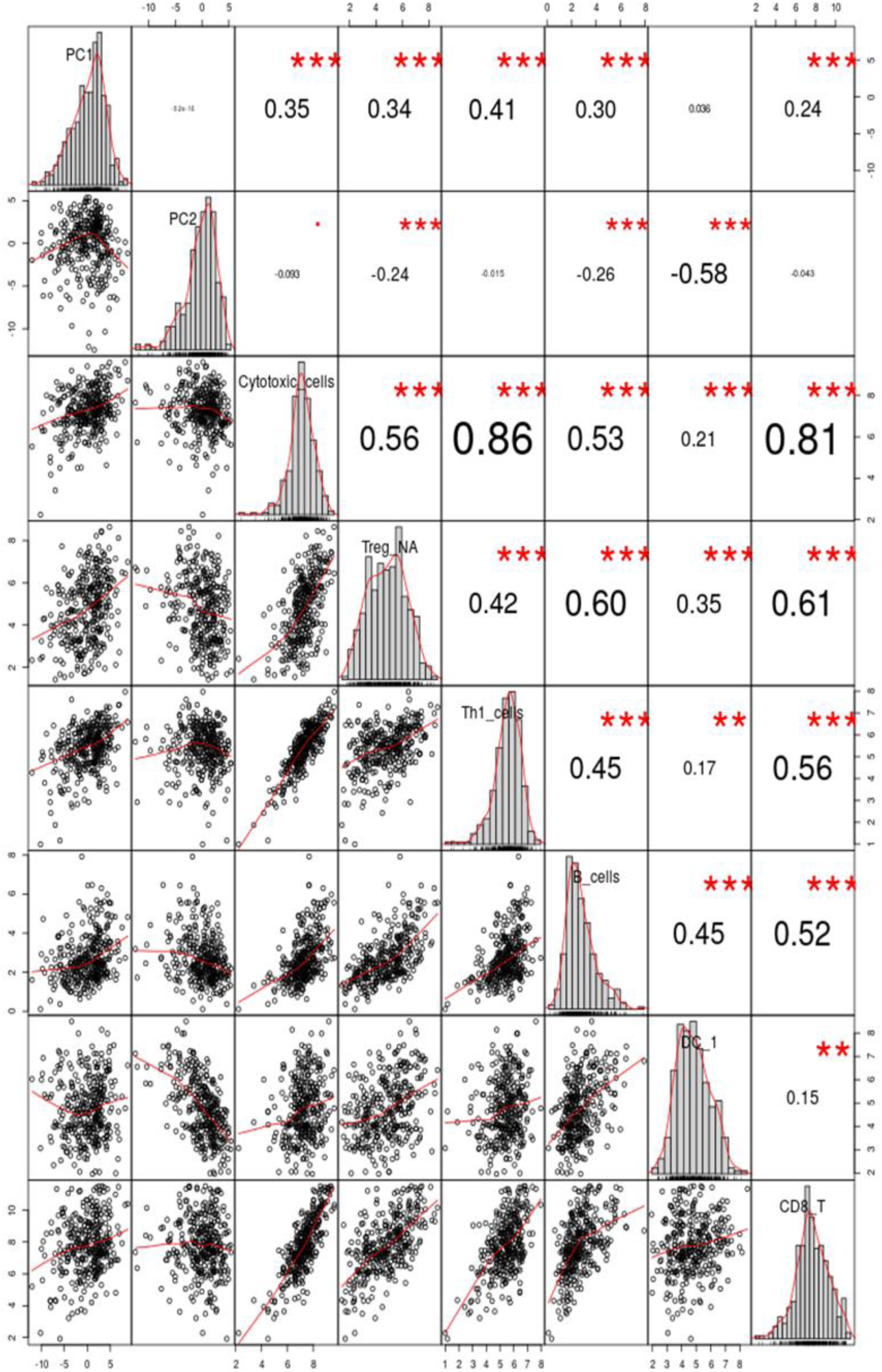
Correlation plot of the PC1, PC2 and the score of TIL.

To further study if PC1 and PC2 were due to the extent of TIL, the TIL values of related cells were added to the coxph model. We hypothesized that if the TIL score of these cells had a strong influence on the PC value, then the coefficient of PC values would no longer be significant after controlling for TIL scores in the cox regression model. We found that PC2 was no longer associated with survival (p>0.5), whereas the P value of PC1 coefficient was still significant (Hazard ratio=1.14; p=0.001). These results indicate that PC2 can be a reflection of the extent of tumor-infiltrating lymphocytes. On the other hand, PC1 is related to survival even after controlling for TIL score. Moreover, transcripts that contribute to PC1 may play an essential role in the development of cancer and need to be further analyzed.

#### (B) Mechanisms of correlated expression of multiple splicing factor genes

To exploit the potential role of PC1 and its top component genes during cancer development, we further investigated the alternative splicing events of PC1’s top component genes. SiR value of these top component genes are correlated to each other. Do the sequences near these splice sites have any common characteristics that make them under the regulation of similar regulatory elements? Since the sequences near these junctions do not have a particularly apparent universal motif, we next investigated whether these sequences are evolutionarily conserved. If they are conservative, then this splicing event may be biologically functional. We downloaded the human Ultraconserved elements (UCE), which refers to the sequence that are identical in at least two species, from the UCbase 2.0 database (Lomonaco et al. 2014). These highly conserved sequences may have critical biological functions. Interestingly, we found that five of the top 10 most contributing genes in PC1 have ultraconserved elements. And four of them have ultraconserved elements near the alternative splice site of their candidate junction (Figure 15-18). Our results indicate that alternative splicing events that produce these candidate junctions are highly conserved and may have particular biological functions. Since most of these PC1 top component genes encode regulatory factors involved in RNA-splicing, their subtle changes in isoform expression may affect the expression of transcripts of the downstream target genes. In summary, the isoform-switch of these top component genes in cancer samples may play an essential role in the development of cancer.

**Figure 15.**
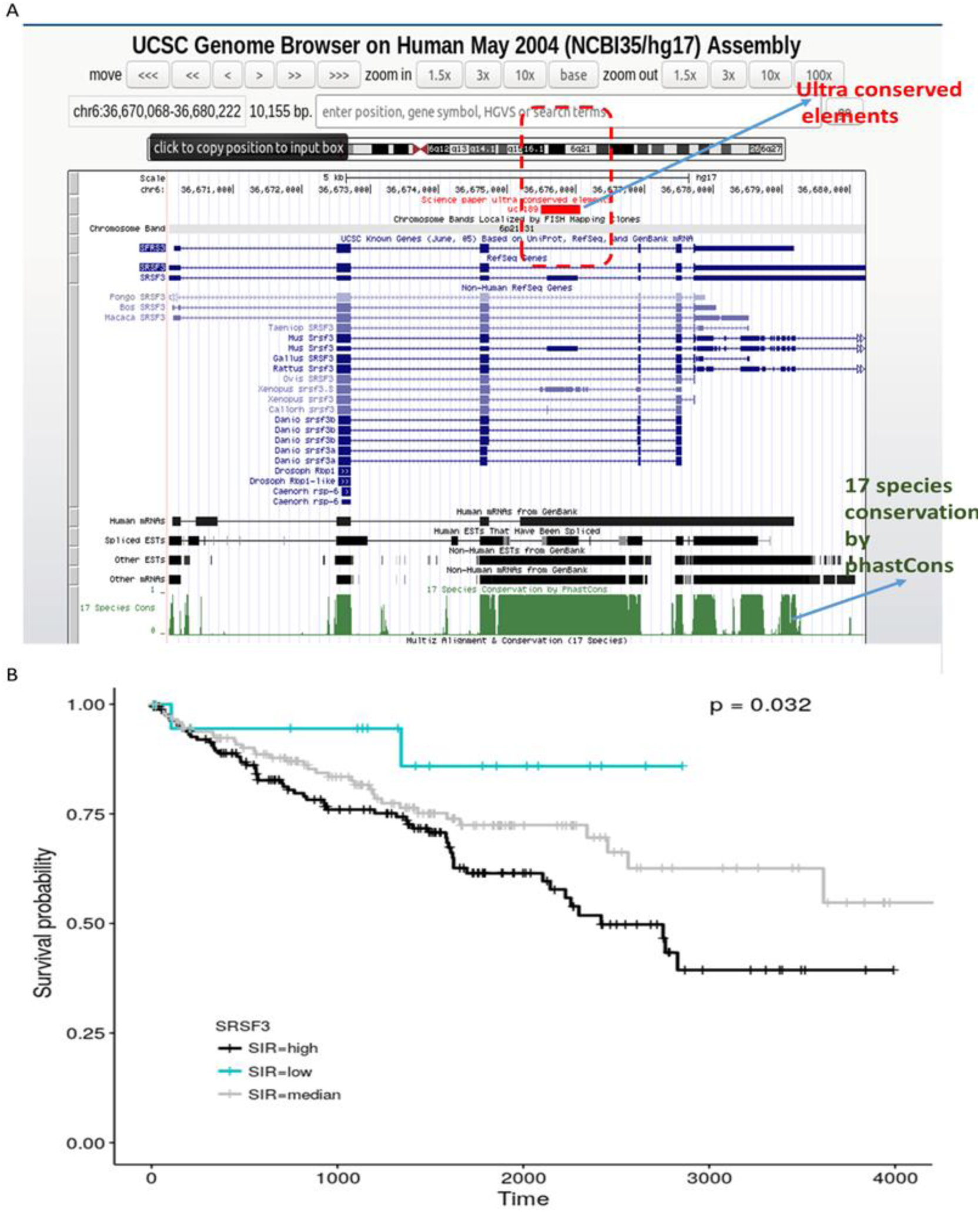
(A) Location of the ultraconserved elements in gene SRSF3 (marked by the red box); (B) Kaplan-Meier plot of the SiR value of SRSF3. High SiR value indicates expression of the non-productive transcript RNA (nt-RNA) including the red box UCE and lead to a premature stop codon in the mRNA of SRSF3.

**Figure 16.**
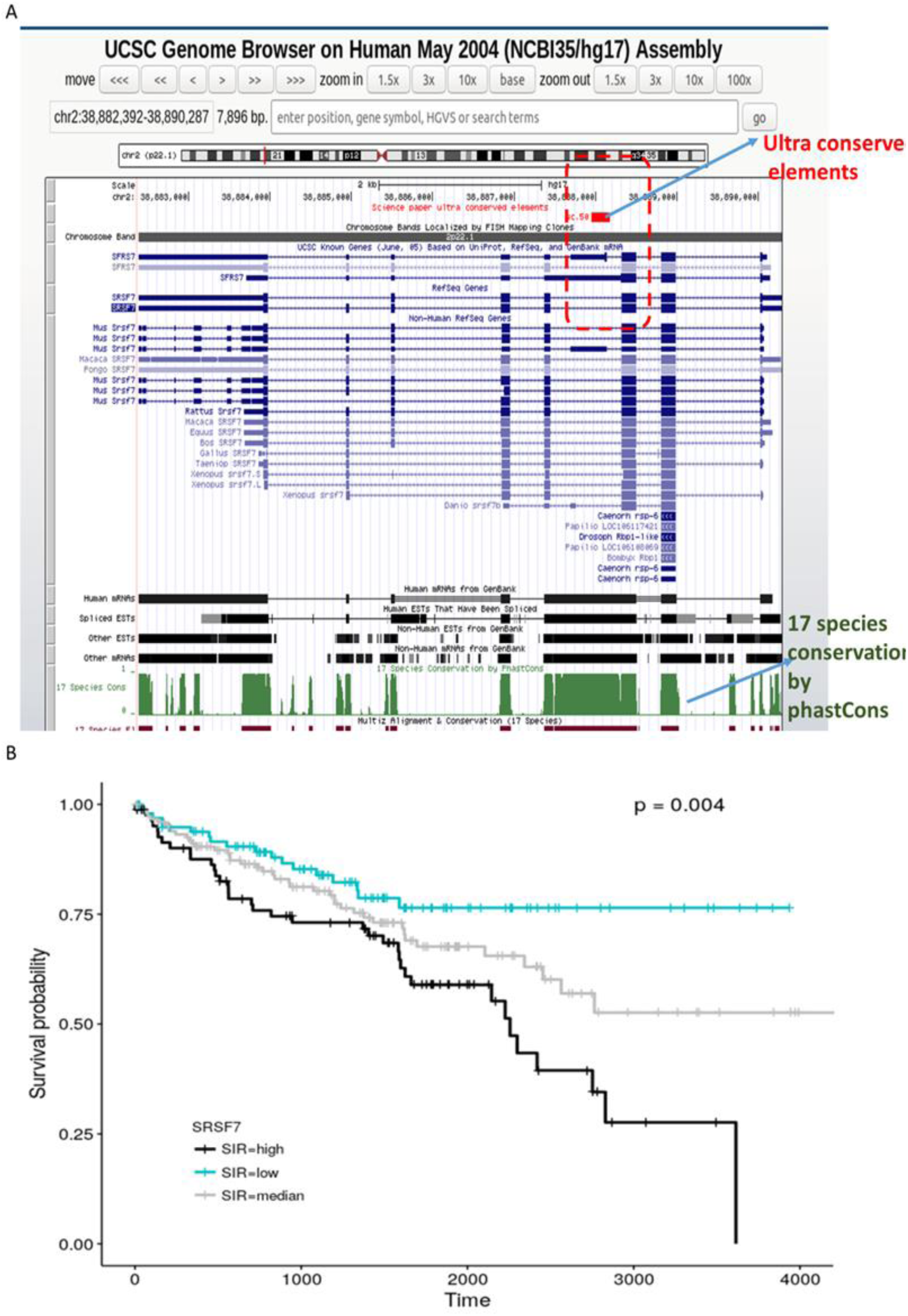
(A) Location of the ultraconserved elements in gene SRSF7 (marked by the red box); (B) Kaplan-Meier plot of the SiR value of SRSF7. High SiR value indicates expression of the non-productive transcript RNA (nt-RNA) including the red box UCE and lead to a premature stop codon in the mRNA of SRSF7.

**Figure 17.**
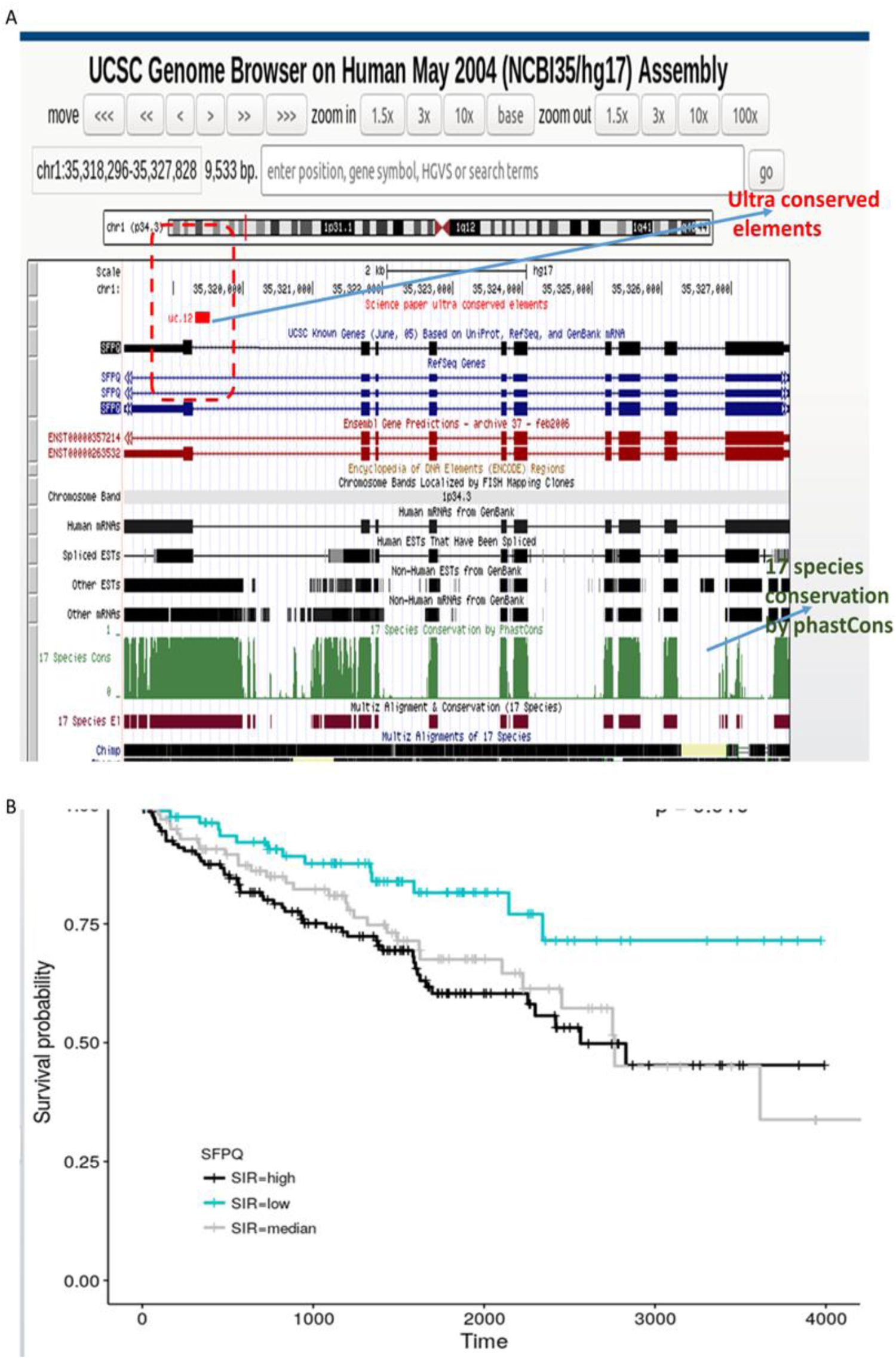
(A) Location of the ultraconserved elements in gene SFPQ (marked by the red box); (B) Kaplan-Meier plot of the SiR value of SFPQ. High SiR value indicates expression of the non-productive transcript RNA (nt-RNA) including the red box UCE and lead to a premature stop codon in the mRNA of SFPQ.

**Figure 18.**
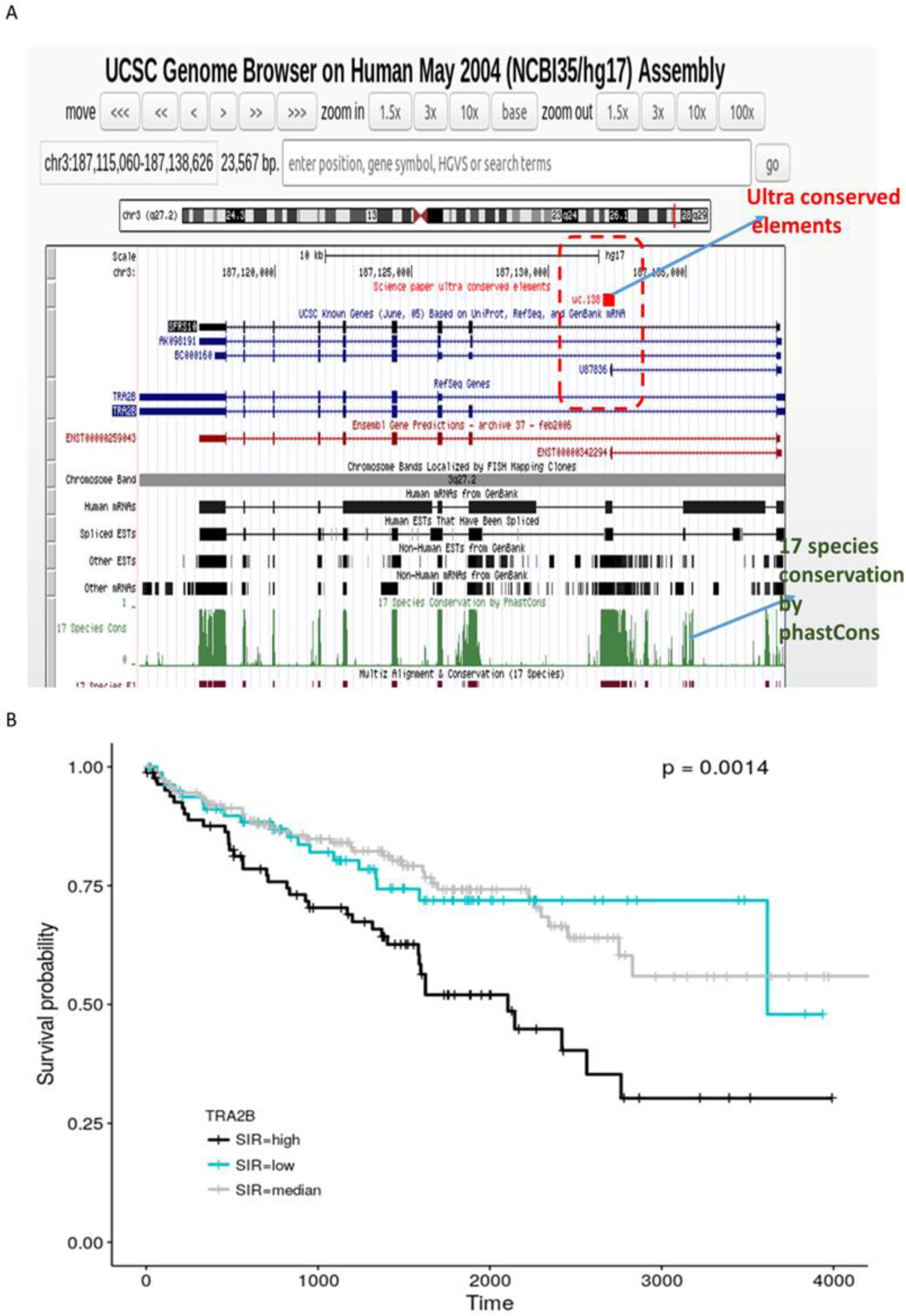
(A) Location of the ultraconserved elements in gene TRA2B (marked by the red box); (B) KM plot of the SiR value of TRA2B. High SiR value indicates expression of the non-productive transcript RNA (nt-RNA) including the red box UCE and lead to a premature stop codon in the mRNA of TRA2B.

#### (C) Mutual regulation mechanism between SRSF3 and CLK1

Among the KIRC PC1 top component gene with highly conserved splice sites, SRSF3’s alternative splicing event not only has an enormous contribution but is ubiquitous in most other cancers. We next wanted to know which genes are involved in or affected by the isoform switch of SRSF3. We only focused on the expression of exon-exon junctions. Junction per million (JPM) was used here, which is normalized values of each junction by dividing by the sequencing depth. The correlation between the JPM value of all the expressed junctions (including NM junction, toxic junction, and novel signature junction) and SiR values of SRSF3 was analyzed. We found that most of the genes most associated with SiR value of SRSF3 are genes of the SR family as well as other genes involved in alternative splicing. This phenomenon may be related to the mechanism by which these splicing regulatory genes interact with each other. At the same time, we also found that the SiR of SRSF3 is also related to the expression values of CLK1 and CLK4 junctions. The CLK family is a class of proteins that phosphorylate other proteins. CLK1 has been shown to be involved in the hyperphosphorylation of SR proteins, and only highly phosphorylated SR proteins are capable of regulating alternative splicing of other genes. Interestingly, CLK1 also has multiple transcripts. Exon4 skipping of CLK1 will produce nonsense transcripts. By comparing the expression value of the signature junction of the transcripts of CLK1 and SRSF3 (Figure 19), we found that the expression of the junction that includes exon4 in isoform of CLK1 (CLK1-I4) and the junction that includes exon4 in isoform of SRSF3 (SRSF3-I4) is highly correlated (r = 0.78). Our results show that the CLK1 protein may further phosphorylate the SRSF3 protein, allowing the SRSF3 protein to function as a splicing promoter which results in the inclusion of the exon4 of itself and the exon4 of CLK1. The CLK1 transcript containing exon4 will be translated into a functional protein, increasing highly phosphorylated SRSF3 protein, while the transcript of SRSF3 contains exon4 is nt-RNA and could not be translated into a functional protein. Such an auto- regulatory feedback mechanism allows the body to control the expression of the SRSF3 protein, thereby also controlling the alternative splicing of its targeted genes. Some articles have experimentally proved that external stimuli, such as increased temperature, will cause transcript conversion of CLK1, while other genes such as SR gene will also undergo transcript conversion (Boutz et al. 2015; Czubaty and Piekiełko-Witkowska 2017). Moreover, inhibition of CLK1 also leads to the isoform-switch in genes such as SRSF3. This result shows that there is indeed a mutual regulation mechanism between SRSF3 and CLK1. Similar results were found by immunohistochemistry study of tumor samples.

**Figure 19.**
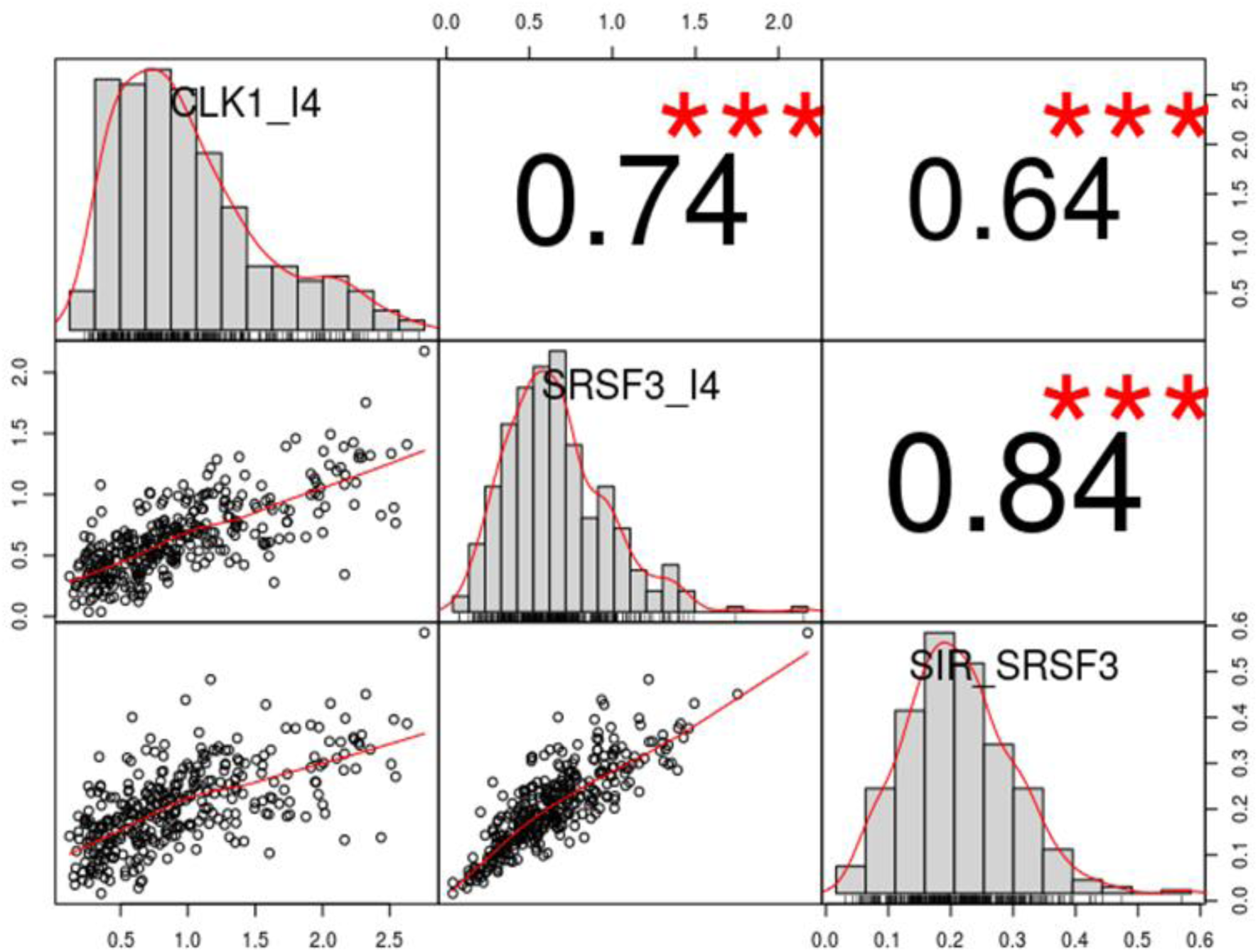
Correlation plot of the SRSF3 and CLK isoforms. Values shown are correlation coefficient.

Next, we primarily clustered candidate junctions that are highly expressed and have variable splicing extents in KIRC cancer samples and obtained two KIRC subtypes associated with survival. Further PCA analysis found that both PC1 and PC2 are related to survival. PC2 is related to TIL, while PC1’s top component gene is most highly conserved and related to RNA splicing. The alternative splicing of SRSF3 and the alternative splicing of CLK1 are mutually adjusted. We also examined the expression and distribution of CLK1 and SRSF3 proteins in renal cell carcinoma (Figure 20). In the same patient, we saw that SRSF3 protein is highly expressed and is mainly in the nucleus, while CLK1 protein has a moderate expression and is primarily in the cytoplasm. Further research may be needed to investigate the regulatory relationship between CLK1 and SRSF3.

**Figure 20.**
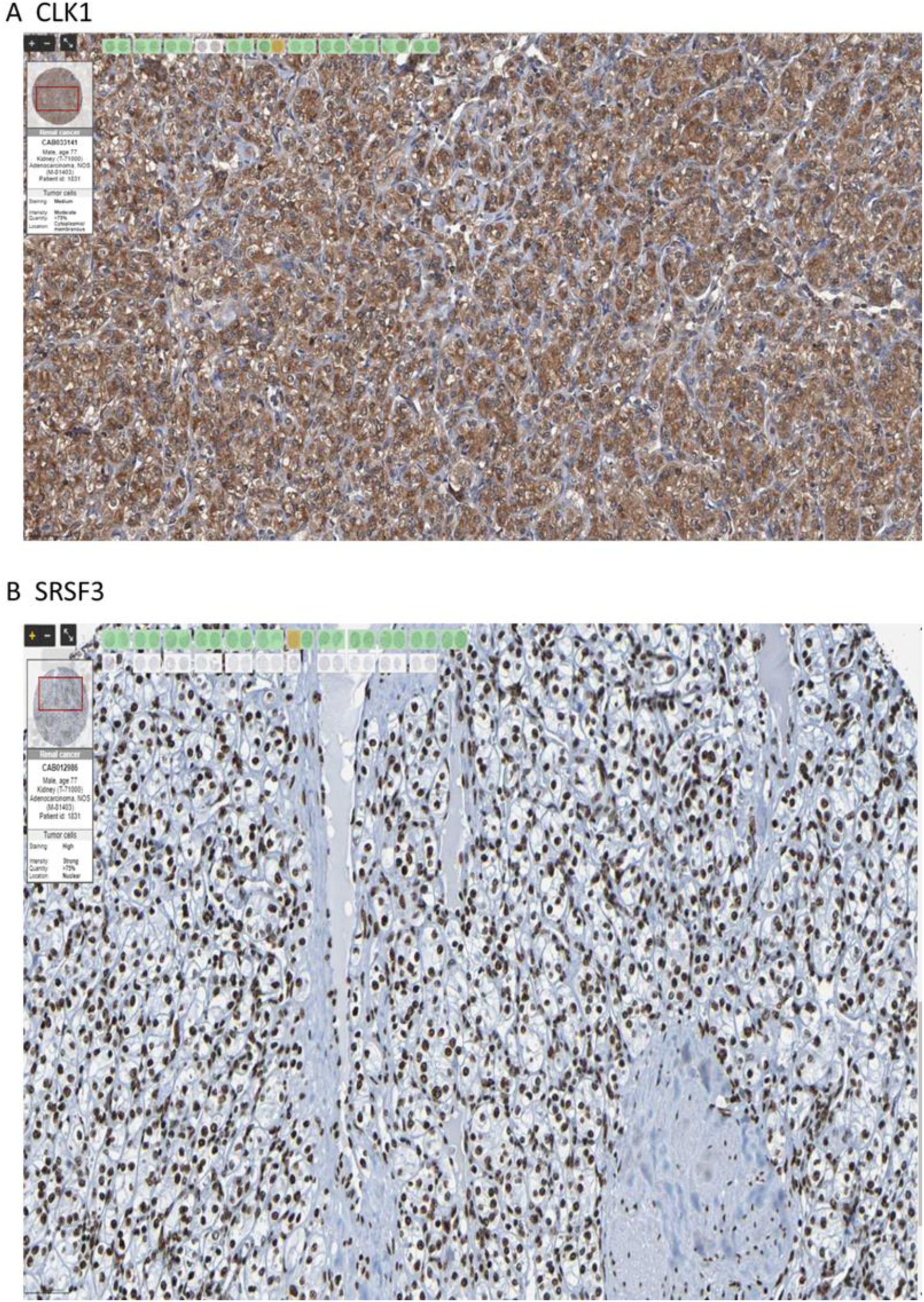
The protein expression of CLK1 and SRSF3 in the same patient with kidney cancer. This was obtained from The Human Protein Atlas (Uhlén et al. 2015). Cancer section of a 77 year old patient shows strong expression of CLK1 and which is associated with nuclear staining of SRSF3.

It was found that unproductive splicing causing accumulation of nt-RNA in some KIRC were associated with poor survival even after corrected for other clinical features. This approach enables identification of novel cancer biomarkers with nt-RNA junctions and provides a mathematical framework to show the dynamics of production of nt-RNA junctions.

## 4 Conclusion

The focus of this study is about a class of transcript isoforms that are defective in term of protein translation, which we called non-translated transcripts (nt-RNA). They are isoform transcripts produced by ASE from the protein-coding gene. However, nt-RNA are produced by specific ASE and leads to frame-shift of the coding sequence or creating a premature stop codon. Most nt-RNA have unique exon-exon junctions that can be used as signature of nt-RNA. Therefore, we can unambiguously identify an nt-RNA transcript by such the signature of these toxic exon-exon junctions. In addition, they can be used as biomarkers which applications are illustrated here.

With the advance of deep sequencing technologies, we investigated the prevalence of alternative splicing and non-translated transcripts (nt-RNAs) in normal and tumor samples across multiple cancer types. Due to the difficulty in quantifying the entire transcript of non-coding RNA, in this study we chose to focus exclusively on the expression level of junctional reads, which span the exon-exon junction of coding or non-coding genes. We profiled the expressed gene junctions of known transcripts and discovered novel signature junctions in RNA-seq data downloaded from the TCGA database.

We found the expression of non-translated transcripts (nt-RNA) is very common and it was produced by most of the expressed protein-coding genes based on the RNA-seq data in discovery datasets. The distribution of the proportional expression of the toxic junction to "normal" splicing junction in different cancer datasets shows that the highest peak was located at around 0.1 in most cancer, which indicates that most of the protein-coding genes only express about 10% or less of the nt-RNA. Interestingly, we have also found that some signature junctions of nt-RNA are expressed in large quantities, accounting for more than 10% of the entire gene junction expression, and some even account for 50%. Those genes may have some potential functions during the tumor progression.

We introduced the Splicing in Ratio (SiR) as a robust method to quantify alternative splicing and identify differentially expressed nt-RNAs and novel junctions between normal and cancer samples. SiR was defined as the raw junction reads count divided by the sum of reads aligned to either the 5’ or 3’ splice site involved in that junction. This value allows us to evaluate the nt-RNA expression ratio more accurately. Through this approach, we identified differentially expressed nt-RNAs and novel junctions in several genes across cancers. For example, we found that ANXA6 and ESYT2 have differentially expressed toxic junctions that potentially play an essential role in cancer development and could be used as potential biomarkers.

Junction profiling using SiR values was successfully applied to identify survival-associated tumor subtypes in KIRC. We selected 173 candidate junctions that are both moderately expressed and showed a variable expression among the KIRC tumor samples. Based on these junctions, we used unsupervised clustering methods and found that these samples assembled into four categories. Two clusters were associated with better survival and two with worse survival, confirming that alternative splicing production data is informative for disease prognosis.

Principal component analysis (PCA) was performed to better understand the signals related to survival. We found that PC1 and PC2 are associated with prognosis and can be used as independent diagnostic biomarkers. The top component genes for PC1 are enriched in gene sets related to RNA splicing, while the top component genes for PC2 are enriched with gene sets which are correlated with the immune cell. We found that PC2 was correlated with tumor-infiltrating lymphocytes (TIL) scores, while PC1 was related to survival even after controlling for TIL score. Moreover, transcripts that contribute to PC1 may play an essential role in the development of cancer.

We investigated the mechanisms of correlated expression of multiple splicing factor genes among PC1’s top component genes and found that their SiR values are correlated to each other. Interestingly, we found that five of the top 10 most contributing genes in PC1 have ultraconserved elements, and four of them have ultraconserved elements near the alternative splice site of their candidate junction. Our results indicate that alternative splicing events that produce these candidate junctions are highly conserved and may have particular biological functions. We also discovered mutual regulation mechanisms between SRSF3 and CLK1, showing that there is indeed a mutual regulation mechanism between these splicing regulatory genes.

In summary, analysis of RNA-seq of cancer transcriptome provides essential information about the gene expression and variation of transcripts. It also provides a means to assess the functional consequence of somatic mutations and characterize novel transcripts. The findings suggest that profiling alternative splicing and non-translated transcripts using metrics like SiR is valuable for cancer research, providing insights into tumor biology, potential biomarkers, and classification. This approach enables identification of novel cancer biomarkers with nt-RNA junctions and provides a mathematical framework to show the dynamics of production of nt- RNA junctions. We found that unproductive splicing causing accumulation of nt-RNA in some KIRC were associated with poor survival even after corrected for other clinical features. These expression profiles can allow researchers to study the expression of toxic junctions or novel signature junctions in or near the genes they are interested in, which could provide a new direction for their research.

## Acknowledgement

The results shown here are in whole or part based upon data generated by the TCGA Research Network: https://www.cancer.gov/tcga.

## Conflict of interest

The authors declare there is no conflict of interest.

## Data availability

The summary statistics of novel toxic junctions and novel junctions are available for download from GitHub. Position of toxic junctions and novel junctions are uploaded to UCSC Genome Browser for easy visualisation. https://github.com/danhuang0909/nt_database

## Notes

https://github.com/danhuang0909/nt_database

### Competing Interest Statement

The authors have declared no competing interest.

## References

1. Arif T, Krelin Y, Shoshan-Barmatz V (2016) Reducing VDAC1 expression induces a non-apoptotic role for pro-apoptotic proteins in cancer cell differentiation. Biochimica et Biophysica Acta (BBA) - Bioenergetics 1857:1228–1242. 10.1016/j.bbabio.2016.04.005

2. Black DL (2003) Mechanisms of Alternative Pre-Messenger RNA Splicing. Annual Review of Biochemistry 72:291–336

3. Boutz PL, Bhutkar A, Sharp PA (2015) Detained introns are a novel, widespread class of post-transcriptionally spliced introns. Genes & Development 29:63–80. 10.1101/gad.247361.114

4. Cotto KC, Feng Y-Y, Ramu A, et al (2023) Integrated analysis of genomic and transcriptomic data for the discovery of splice-associated variants in cancer. Nature Communications 14:1589. 10.1038/s41467-023-37266-6

5. Czubaty A, Piekiełko-Witkowska A (2017) Protein kinases that phosphorylate splicing factors: Roles in cancer development, progression and possible therapeutic options. The International Journal of Biochemistry & Cell Biology 91:102–115. 10.1016/j.biocel.2017.05.024

6. Danaher P, Warren S, Dennis L, et al (2017) Gene expression markers of Tumor Infiltrating Leukocytes. J Immunother Cancer 5:18. 10.1186/s40425-017-0215-8

7. de Muga SV, Timpson P, Cubells L, et al (2009) Annexin A6 inhibits Ras signalling in breast cancer cells. Oncogene 28:363–377. 10.1038/onc.2008.386

8. Fair B, Buen Abad Najar CF, Zhao J, et al (2024) Global impact of unproductive splicing on human gene expression. Nature Genetics 56:1851–1861. 10.1038/s41588-024-01872-x

9. Grewal T, Koese M, Rentero C, Enrich C (2010) Annexin A6-regulator of the EGFR/Ras signalling pathway and cholesterol homeostasis. The International Journal of Biochemistry & Cell Biology 42:580–584. 10.1016/j.biocel.2009.12.020

10. Jean S, Mikryukov A, Tremblay MG, et al (2010) Extended-Synaptotagmin-2 Mediates FGF Receptor Endocytosis and ERK Activation In Vivo. Developmental Cell 19:426–439. 10.1016/j.devcel.2010.08.007

11. Kahles A, Lehmann K-V, Toussaint NC, et al (2018) Comprehensive Analysis of Alternative Splicing Across Tumors from 8,705 Patients. Cancer Cell 34:211–224.e6. 10.1016/j.ccell.2018.07.001

12. Koese M, Rentero C, Kota BP, et al (2013) Annexin A6 is a scaffold for PKCα to promote EGFR inactivation. Oncogene 32:2858–2872. 10.1038/onc.2012.303

13. Krawczak M, Thomas NST, Hundrieser B, et al (2007) Single base-pair substitutions in exon–intron junctions of human genes: nature, distribution, and consequences for mRNA splicing. Human Mutation 28:150–158. 10.1002/humu.20400

14. Li B, Severson E, Pignon J-C, et al (2016) Comprehensive analyses of tumor immunity: implications for cancer immunotherapy. Genome Biology 17:174. 10.1186/s13059-016-1028-7

15. Li H, Handsaker B, Wysoker A, et al (2009) The Sequence Alignment/Map format and SAMtools. Bioinformatics 25:2078–2079. 10.1093/bioinformatics/btp352

16. Liberzon A, Subramanian A, Pinchback R, et al (2011) Molecular signatures database (MSigDB) 3.0. Bioinformatics 27:1739–1740. 10.1093/bioinformatics/btr260

17. Lomonaco V, Martoglia R, Mandreoli F, et al (2014) UCbase 2.0: ultraconserved sequences database (2014 update). Database 2014:bau062. 10.1093/database/bau062

18. Pan Q, Shai O, Lee LJ, et al (2008) Deep surveying of alternative splicing complexity in the human transcriptome by high-throughput sequencing. Nature Genetics 40:1413–1415. 10.1038/ng.259

19. Qi H, Liu S, Guo C, et al (2015) Role of annexin A6 in cancer (Review). Oncol Lett 10:1947–1952. 10.3892/ol.2015.3498

20. Quinlan AR, Hall IM (2010) BEDTools: a flexible suite of utilities for comparing genomic features. Bioinformatics 26:841–842. 10.1093/bioinformatics/btq033

21. Rini BI, Campbell SC, Escudier B (2009) Renal cell carcinoma. The Lancet 373:1119–1132. 10.1016/S0140-6736(09)60229-4

22. Sakwe AM, Koumangoye R, Guillory B, Ochieng J (2011) Annexin A6 contributes to the invasiveness of breast carcinoma cells by influencing the organization and localization of functional focal adhesions. Experimental Cell Research 317:823–837. 10.1016/j.yexcr.2010.12.008

23. Seiler M, Peng S, Agrawal AA, et al (2018) Somatic Mutational Landscape of Splicing Factor Genes and Their Functional Consequences across 33 Cancer Types. Cell Reports 23:282–296.e4. 10.1016/j.celrep.2018.01.088

24. Smith CWJ, Valcárcel J (2000) Alternative pre-mRNA splicing: the logic of combinatorial control. Trends in Biochemical Sciences 25:381–388. 10.1016/S0968-0004(00)01604-2

25. Strzelecka-Kiliszek A, Buszewska ME, Podszywalow-Bartnicka P, et al (2008) Calcium- and pH-dependent localization of annexin A6 isoforms in Balb/3T3 fibroblasts reflecting their potential participation in vesicular transport. Journal of Cellular Biochemistry 104:418–434. 10.1002/jcb.21632

26. Thierry-Mieg D, Thierry-Mieg J (2006) AceView: a comprehensive cDNA-supported gene and transcripts annotation. Genome Biology 7:S12. 10.1186/gb-2006-7-s1-s12

27. Tremblay MG, Herdman C, Guillou F, et al (2015) Extended Synaptotagmin Interaction with the Fibroblast Growth Factor Receptor Depends on Receptor Conformation, Not Catalytic Activity *. Journal of Biological Chemistry 290:16142–16156. 10.1074/jbc.M115.656918

28. Uhlén M, Fagerberg L, Hallström BM, et al (2015) Tissue-based map of the human proteome. Science 347:1260419. 10.1126/science.1260419

29. Varn FS, Tafe,Laura J., Amos,Christopher I., and Cheng C (2018) Computational immune profiling in lung adenocarcinoma reveals reproducible prognostic associations with implications for immunotherapy. OncoImmunology 7:e1431084. 10.1080/2162402X.2018.1431084

30. Vitting-Seerup K, Sandelin A (2017) The Landscape of Isoform Switches in Human Cancers. Molecular Cancer Research 15:1206–1220. 10.1158/1541-7786.MCR-16-0459

31. Wang ET, Sandberg R, Luo S, et al (2008) Alternative isoform regulation in human tissue transcriptomes. Nature 456:470–476. 10.1038/nature07509

32. Wang Z, Burge CB (2008) Splicing regulation: From a parts list of regulatory elements to an integrated splicing code. RNA 14:802–813. 10.1261/rna.876308

33. Wang Z, Gerstein M, Snyder M (2009) RNA-Seq: a revolutionary tool for transcriptomics. Nature Reviews Genetics 10:57–63. 10.1038/nrg2484

34. Zhang Z, Pal S, Bi Y, et al (2013) Isoform level expression profiles provide better cancer signatures than gene level expression profiles. Genome Medicine 5:33. 10.1186/gm437

35. Zhao W, Hoadley KA, Parker JS, Perou CM (2016) Identification of mRNA isoform switching in breast cancer. BMC Genomics 17:181. 10.1186/s12864-016-2521-9

