## Supplementary material for "A map of Non-translated RNA (nt-RNA) junctions in cancer genomes: a database resource of unproductive splicing"

Supplementary document

(1) Explanation of the toxic junctions is the unique tag of many nt-RNA causing frame-shift of ORF

A

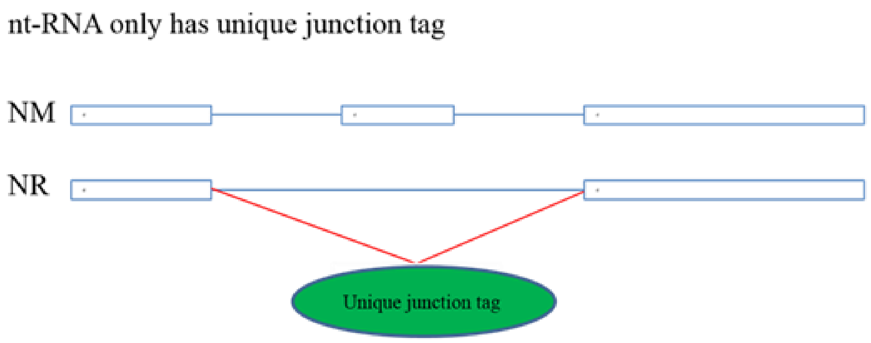


B

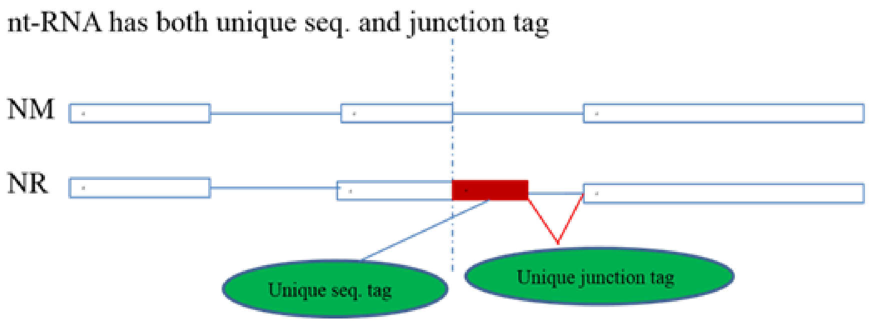


A and B show 2 examples of ASE leading to a frame-shift ORF. NM represents the canonical transcripts. NR represents nt-RNA. In example A, there is a unique junction found only in NR nt-RNA transcript. In example B, there is a short unique sequence tag in addition to the unique junction tag. Therefore, unique junction tags are generated in most of the nt-RNA when compared to the canonical transcript. From the genome database, these tags retrieved from known annotated nt-RNA are referred to as NR junctions. Other unannotated nt-RNA or junctions associated with nt-RNA are called novel junctions. They are called toxic junctions.

**Both types of toxic junctions are shown in the UCSC browser.**

### (2) Prioritise annotated transcripts (NM sequences) to obtain the frame information to identify toxic junctions.

The algorithm for assigning frame information to novel junctions is explained here.

(A) First, a model of an annotated transcript aligning to both sides of the new junction Is selected by a priority scoring method.
The transcript with the top sum of 3 score is selected, e.g. the highlight one with a sum of score = 12, 5+4+3

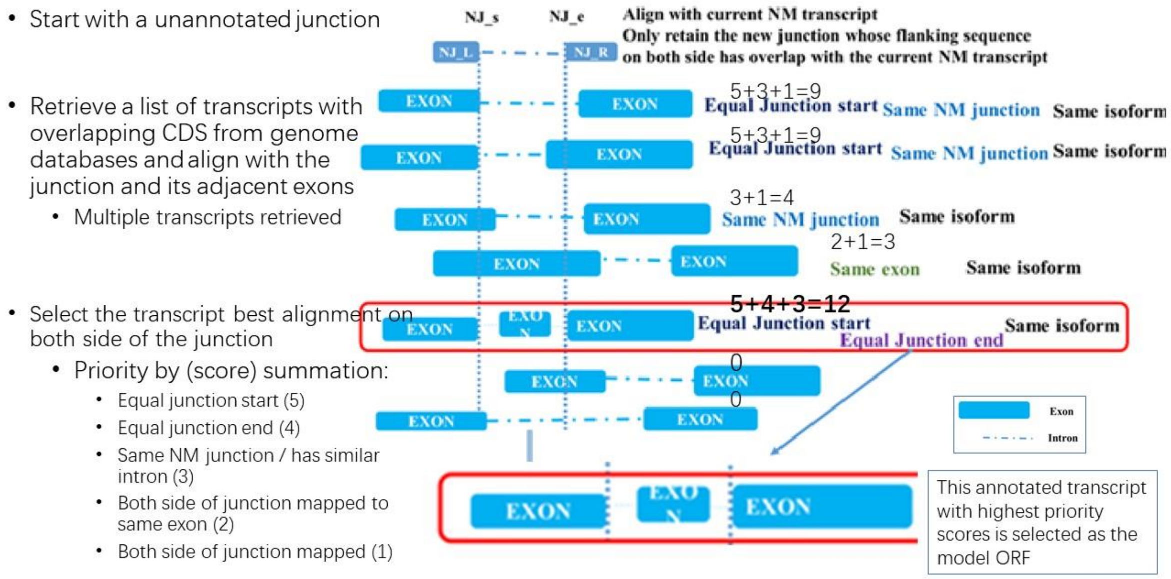


(B) Then the difference (d) of intron bases (length) is calculated to determine if it is a multiple of 3.

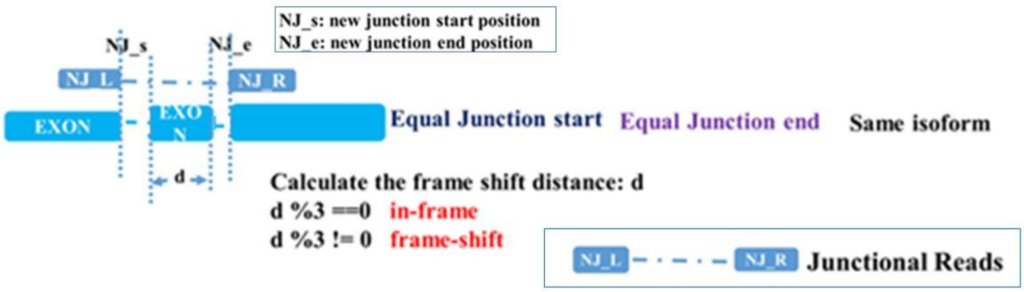


- Novel junctional reads are aligned to the selected annotated model transcript which provide the frame information of ORF.
- The frame shift distance (d) is calculated by difference between intron bases (length) between the novel junction and the model transcript between the ends of the intron derived by novel junction.
- When d is not a multiple of 3 bp, it indicates a frame shift due to this ASE, the novel junction is a toxic junction.
